# Exploring the Regulation of Cdc42 Stability and Turnover in Yeast

**DOI:** 10.1101/2022.09.30.510332

**Authors:** Beatriz González, Martí Aldea, Paul J. Cullen

**Affiliations:** Department of Biological Sciences, State University of New York at Buffalo; Molecular Biology Institute of Barcelona (IBMB), CSIC, Barcelona, Spain

**Author notes:** Corresponding author: Paul J. Cullen, 532 Cooke Hall, Department of Biological Sciences, State University of New York at Buffalo, Buffalo, NY 14260-1300.

**Keywords:** Temperature, protein trafficking, ESCRT, sexual selection, tradeoffs, quality control, protein aggregation, aging

## Abstract

Rho GTPases govern many cellular processes, including actin cytoskeleton dynamics and signal transduction pathways. Rho GTPase levels can be regulated by stability and turnover, yet many aspects of this type of regulation remain largely unexplored. We report here a new environmental stress, high temperature (37°C), that stimulates yeast Cdc42p turnover to impact its biological functions. At 37°C, Cdc42p turnover required the NEDD4 ubiquitin ligase Rsp5p and HSP40/HSP70 chaperones. Specific lysine residues promoted Cdc42p degradation at 37°C [K166; and residues in the Poly-Basic (PB) domain: K183, K184, K186, K187], which occurred in both the 26S proteosome and ESCRT-to-vacuole pathway. Degradation of Cdc42p at 37°C reduced the sensitivity to mating pheromone, demonstrating biological role for Cdc42p turnover in this context. Stabilization of Cdc42p at high temperatures restored pheromone sensitivity but caused growth and polarity defects, suggesting a tradeoff between sexual propagation and cellular fitness. One lysine residue (K16) in the P-loop of the protein was critical for stability. Overproduction of the protein, expression of Cdc42p^K16R^ in a mutant where the protein accumulates, and other types of proteostatic stress led to the formation of Cdc42p aggregates in aging mother cells. These new aspects of Cdc42p protein quality control may extend to other members of the Rho GTPase family of proteins.

**Summary statement:** Rho GTPases regulate cell polarity and signaling (e.g. MAPK) pathways. Here, we discovered that yeast Cdc42p is targeted for degradation at 37°C by a NEDD4 ubiquitin ligase and HSP40 and HSP70 chaperones through lysine residues in the C-terminus of the protein. At 37°C, Cdc42p was degraded both by the 26S proteasome and in an ESCRT-dependent manner in the vacuole. Preventing Cdc42p turnover at 37°C resulted in improved mating sensitivity but also viability and polarity defects, suggesting a tradeoff between sexual responses and fitness. In addition, one residue (K16) was critical for Cdc42p stability. Cdc42p^K16R^ formed aggregates in aging mother cells, and aggregates were also observed in cells undergoing proteostatic stress. Protein quality control regulation of a Rho-type GTPase therefore has ramification in the regulation of cellular responses, evolutionary tradeoffs, and protein aggregation in ways that might impact aging.

**HIGHLIGHTS:**

i. **High temperatures** (37°C) induce turnover of the Rho GTPase Cdc42p
ii. Turnover of Cdc42p at 37°C requires the **HSP40/HSP70** proteins and the NEDD4-type **E3 ubiquitin ligase Rsp5p.**
iii. **K166** and four lysines at the extreme C-terminus [poly-basic (PB: **K183, K184, K186, K187**] promote **turnover of Cdc42p** at 37°C
iv. Cdc42p is degraded at 37°C by the proteosome and the ESCRT-to-vacuole pathways.
v. GTP-Cdc42p does not accumulate in ESCRT mutants and is not turned over in the vacuole.
vi. Turnover of Cdc42p at 37°C inhibits sensitivity to mating pheromone Preventing Cdc42p turnover restores pheromone sensitivity at the cost of cell viability and proper cell polarity. These results reveal a tradeoff between sexual responses and overall cellular fitness.
vii. An internal lysine residue (K16) is required for Cdc42p stability.
  i. verproduction of the protein, or accumulation of Cdc42p^K16R^ in certain mutants induces **protein aggregation** in aging mother cells.

## INTRODUCTION

Small GTPases (21-25 kDa) in the Ras (Rat Sarcoma Virus) superfamily of monomeric GTP binding proteins control a diverse range of cellular processes, including cell invasion, migration, proliferation and survival. A prominent group in this family are Rho (Ras homology) GTPases (e.g. Cdc42, Rac1, and RhoA), which mainly function to regulate cytoskeletal dynamics and signal transduction pathways (Etienne-Manneville and Hall, 2002; Ridley, 2011). Rho GTPases function as molecular switches, which are active when bound to GTP and inactive when bound to GDP. The active or GTP-bound state leads to interaction with a wide variety of effector proteins, including formins that regulate the actin cytoskeleton, and protein kinases that stimulate signaling pathways. These effector proteins can modulate the shape and migration of cells, as well as cell cycle progression, gene expression, apoptosis and survival (Coleman et al., 2004; Coso et al., 1995; Lai et al., 2002; Lawson and Ridley, 2018; Prudnikova et al., 2015). Guanine nucleotide exchange factors (GEFs) catalyze the exchange of GDP for GTP to activate GTPases, and GTPase activating proteins (GAPs) promote the intrinsic GTP hydrolysis of GTPases, to promote cycling to their inactive conformation (Bos et al., 2007). Guanine nucleotide dissociation inhibitors (GDIs) extract Rho GTPases from membranes thereby preventing the activation by GEFs (Golding et al., 2019).

On top of this core layer of regulation are post-translational modifications (PTMs), which also control the spatiotemporal regulation and activity of Rho GTPases. One type of regulation impacts Rho GTPase stability, which is mainly controlled by ubiquitination (Deng and Huang, 2014; Guo and Rahmouni, 2019; Majolée et al., 2021; Vanneste et al., 2020; Wei et al., 2013). Indeed, many G-proteins, including RAS (Abe et al., 2020; Bigenzahn et al., 2018; Campbell and Philips, 2021), Ras-like proteins (Cuevas-Navarro et al., 2022), and heterotrimeric G-proteins that function with seven-transmembrane receptors (Dohlman and Campbell, 2019; Wang et al., 2005) are similarly regulated. Rho GTPases are also regulated by other PTMs, including lipid modifications (Michaelson et al., 2001; Navarro-Lérida et al., 2012), phosphorylation (Chang et al., 2011; Cho et al., 2018; Tkachenko et al., 2011), SUMOylation (Yue et al., 2017), and oxidation (Mitchell et al., 2013), which can modulate GTPase localization, activity and function in specific contexts.

Although many aspects of Rho GTPase stability remain relatively unexplored, it is clear that mis-regulation of protein levels can have a dramatic impact in several human diseases including cancer (Clayton and Ridley, 2020; Goka and Lippman, 2015; Haga and Ridley, 2016; Li et al., 2016) and neurodegenerative disorders (Arrazola Sastre et al., 2020). In particular, CDC42 activity has been proposed to impact cell migration and invasion during malignant transformation (Reymond et al., 2012; Svensmark and Brakebusch, 2019; Zhang et al., 2019), as well as cellular aging (Florian et al., 2012; Geiger and Zheng, 2013). Thus, revealing the molecular mechanisms that regulate CDC42 stability and turnover might provide new insights into human diseases and aging.

Proteins in the cell can be turned over after ubiquitination by two major pathways. One pathway requires the 26S proteosome (Amm et al., 2014; Finley et al., 2012). Another pathway involves the delivery of proteins by intracellular trafficking to the lysosome or vacuole in yeast (Settembre et al., 2013; Wang et al., 2018). <u>E</u>ndosomal <u>s</u>orting <u>c</u>omplexes required for transport (ESCRT) direct cargos internalized from the plasma membrane to the inside of endosomes through invagination, resulting in their localization in the prevacuolar compartment or multivesicular bodies [MVBs, (Katzmann et al., 2001; Remec Pavlin and Hurley, 2020)]. In addition to protein trafficking, ESCRT has recently been described to be required for sealing of the autophagosome membrane during autophagy (Zhou et al., 2019), and is also involved in a wide range of cellular processes, including cytokinesis, neurogenesis, and nuclear envelope regulation (Lu and Drubin, 2020; Vietri et al., 2020). Moreover, proteins can also be directly targeted to the vacuole (Wang et al., 2018) and turned over by autophagy (Farré and Subramani, 2016; Yim and Mizushima, 2020). The yeast E3 Rsp5p is involved several biological processes by controlling the turnover of many proteins which can be degraded by the 26S proteosome (Brückner et al., 2011; Chernova et al., 2011; Fang et al., 2014; Sommer et al., 2014), including GTP-Cdc42p (González and Cullen, IN PRESS). More commonly, Rsp5p controls the turnover of plasma membrane proteins in the vacuole (Hoshikawa et al., 2003; Lin et al., 2008; Pizzirusso and Chang, 2004; Rotin et al., 2000; Yoshida et al., 2012). Recently, Rsp5p has been found to mediate the ubiquitination of Heat Shock Proteins (HSP), including Hsp70-Ssb1p, Hsp82p and Hsp104p chaperones in a proteasome-dependent manner (Wang et al., 2021). HSP proteins encompass a large family of evolutionary conserved molecular chaperones which mainly control protein homeostasis, based on their role in regulating protein maturation, trafficking, re-folding and degradation (Balchin et al., 2016; Rosenzweig et al., 2019). Based on these several functions, HSP proteins play a critical role in several human diseases (Albakova et al., 2022; Ghadban et al., 2017; Gorenberg and Chandra, 2017; Horianopoulos and Kronstad, 2021; Klaips et al., 2018).

Cdc42p is highly conserved from yeast to humans (81% sequence identity) (Kozminski et al., 2000), which has led to the use of model systems to understand Cdc42p regulation in higher organisms. In the budding yeast *Saccharomyces* cerevisiae, Cdc42p is an essential protein and the principal regulator of cell polarity, during budding, mating and filamentous growth (Bi and Park, 2012; Johnson, 1999; Miller et al., 2020; Park and Bi, 2007). By interaction with effectors, including the formin Bni1p (Evangelista et al., 1997; Evangelista et al., 2002), Gic proteins (Brown et al., 1997; Kawasaki et al., 2003), and p21-activated kinase PAKs (Cvrcková et al., 1995; Evangelista et al., 2002; Gulli et al., 2000), Cdc42p controls the establishment of cell polarity and the formation of a new bud (Martin, 2015; Miller et al., 2020; Moran et al., 2019; Slaughter et al., 2009). Cdc42p can also bind to Ste20p to regulate <u>M</u>itogen-<u>A</u>ctivated <u>P</u>rotein <u>K</u>inase (MAPK) pathways (Cvrcková et al., 1995; Peter et al., 1996; Saito, 2010; Simon et al., 1995). Three Cdc42p-dependent MAPK pathways regulate different cellular responses, including filamentous growth (fMAPK) (Cullen and Sprague, 2012), mating (Good et al., 2009; Sprague et al., 1983), and the response to osmotic stress (HOG) (Saito and Posas, 2012). Understanding how Cdc42 and other common proteins function in a pathway-specific response has been intensively explored (Bao et al., 2004; Basu et al., 2020; Boulter et al., 2010; Patterson et al., 2021; Prabhakar et al., 2021; Saito, 2010; Van Drogen et al., 2020; Vázquez-Ibarra et al., 2020). Cdc42p also regulates exocytosis (Adamo et al., 2001), endocytosis (Aguilar et al., 2006), vacuolar functions (Jones et al., 2010), and nuclear membrane disassembly and migration (Lu and Drubin, 2020). However, many questions about the regulation of Cdc42p stability and its impact on biological functions remain unknown.

We previously showed that heat shock proteins (HSPs) of the HSP40/HSP70 family direct GTP-bound Cdc42p (called GTP-Cdc42p in this study) to the E3 ubiquitin ligase Rsp5p for turnover in the proteosome (González and Cullen, IN PRESS). This discovery prompted examination of other aspects of Cdc42p stability and turnover. HSPs control protein folding and turnover at high temperatures, which suggested that temperature may play a role in Cdc42p turnover regulation. In line with this possibility, we show here that Cdc42p is also turned over at 37°C, which required the same HSP40/HSP70 and Rsp5p proteins. We also identified lysine residues specifically required for turnover of Cdc42p at 37°C. Moreover, whereas GTP-Cdc42p was turned over in the proteosome, at 37°C, Cdc42p was turned over by the proteosome and in the ESCRT-to-vacuole pathway, which represents the first example for turnover of a Rho-type GTPase in this compartment. Turnover of Cdc42p at 37°C reduced the sensitivity to mating pheromone, which indicates that that the turnover of Cdc42p is functionally relevant in this context. Interfering with Cdc42p turnover at 37°C restored pheromone sensitivity but also led to defects in viability and cell polarity, suggesting that Cdc42p degradation at 37°C is required for proper cell function. Finally, we identified one residue (K16) that was critical for the stability of the protein. Accumulation of Cdc42p^K16R^ (or overproduction of the protein) led to the formation of protein aggregates. Our findings provide new insights into the molecular basis of Cdc42p stability and turnover, and their functional consequences on biological activities, ranging to sexual selection tradeoffs to aging. These discoveries may extend to the regulation of Rho GTPases in many settings.

## RESULTS

### Cdc42p is Degraded at High Temperatures (37°C) in an HSP40/HSP70- and Rsp5p-Dependent Manner

Cdc42p protein levels were examined by growth of cells in different environments. Immunoblot analysis using antibodies against the Cdc42p protein showed reduced protein levels during growth at 37°C (Fig. 1A). Immunoblot analysis using anti-GFP antibodies showed that a functional GFP-Cdc42p fusion protein showed a similar pattern (*Fig. S1A*). Cdc42p was not turned over by incubation of cells at 30°C (Fig. 1B; for quantification see Fig. 1C). The protein kinase C (PKC) pathway regulates the response to temperature and other environmental stresses (Heinisch and Rodicio, 2018; Kamada et al., 1995) but was not required for turnover of Cdc42p at 37°C (*Fig. S1B*). Different strain backgrounds showed different rates of turnover of Cdc42p at 37°C (S288c showed slower turnover kinetics than ∑1278b, which was focused on here). The difference in turnover rates could be because the strains have different degrees of thermotolerance, although we have not explored this possibility.

**Figure 1.**
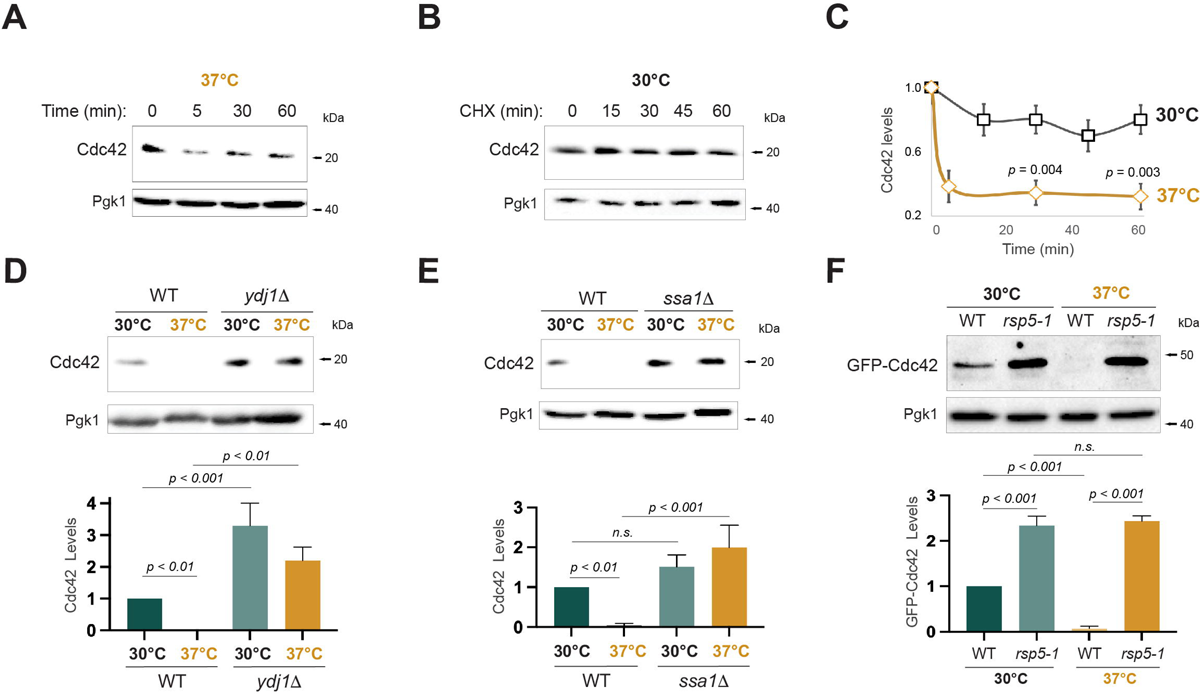
Degradation of Cdc42p is induced at 37°C in a chaperone- and Rsp5p-dependent manner. **A)** Cdc42p levels in wild-type (WT, ∑1278b, PC538) cells incubated for 4h at 30°C and shifted to 37°C for the indicated time points. Anti-Cdc42p and anti-Pgk1 antibodies were used. **B)** Cdc42p levels in wild-type (WT) cells grown in media containing 25 µg/ml of cycloheximide at 30°C and analyzed by immunoblotting at the indicated time points. See Fig. 1A for details. **C)** Relative protein quantification of Cdc42p at 30°C (panel 1B, black) and at 37°C (panel 1A, yellow) to account for the turnover rates. Error bars represent the S.D from two biological replicates. Data were analyzed by one-way ANOVA and Tukey’s multiple comparison test. **D)** Cdc42p levels in wild-type (WT, S288C, PC986) cells and the *ydj1*Δ (PC7657) mutant grown at 30°C for 5 h (30°C) and shifted to 37°C for 2 h (37°C). Anti-Cdc42p and anti-Pgk1 antibodies were used. Bottom, graph represents quantification of the relative levels of Cdc42 compared to Pgk1 from three biological replicates, n = 3. Data were analyzed by one-way ANOVA followed by a Tukey’s multiple comparison test. **E)** Cdc42p levels in wild-type (WT, ∑1278b, PC6016) cells and the *ssa1*Δ (PC7700) mutant grown at 30°C for 5 h (30°C) or shifted to 37°C for 2 h (37°C). See Fig. 1D for details. **F)** Levels of GFP-Cdc42p (PC6454) expressed in wild-type (WT, S288C, PC3288) and *rsp5-1* (PC3290) cells incubated 5 h at 30°C (30°C) and 2h at 37°C (37°C). See Fig. 1D for details. Same results were shown in Gonzalez and Cullen, IN PRESS.

HSP chaperones are upregulated during heat stress to promote protein folding and turnover (Farhan et al., 2021; Lindquist and Craig, 1988). These molecular chaperones fold proteins in an ATP-dependent manner, and direct mis-folded proteins to protein degradation machinery (Balchin et al., 2016; Hartl et al., 2011). We previously showed that the HSP40-type chaperone Ydj1p (Lu and Cyr, 1998; Reidy et al., 2014) is required for turnover of GTP-Cdc42p (Gonzalez and Cullen, IN PRESS). At 37°C, Cdc42p turnover at 37°C was also dependent on Ydj1p (Fig. 1D). HSP40 proteins function as a co-chaperones for HSP70 proteins by controlling the rate of ATP hydrolysis and directing HSP70s to function in a variety of processes (Craig and Marszalek, 2017). At 37°C, an HSP70 chaperone, Ssa1p, was also required for Cdc42p turnover (Fig. 1E). HSP40 proteins can target client proteins for degradation through the NEDD4-type ubiquitin ligase Rsp5p (Fang et al., 2014). We previously showed that Rsp5p is required for degradation of GTP-Cdc42p (Gonzalez and Cullen, IN PRESS). Immunoblot analysis showed that GFP-Cdc42p was also stabilized in the *rsp5-1* mutant at 37°C (Fig. 1F). High molecular weight (HMW) products, which are indicative of poly-ubiquitination (Grice and Nathan, 2016), also accumulated at 37°C in a Rsp5p-dependent manner (*Fig. S1C*), suggesting that Rsp5p is required for poly-ubiquitination of Cdc42p at 37°C. Therefore, high temperature (37°C) is a new stimulus that induces Cdc42p degradation in a HSP40/HSP70- and Rsp5p-dependent manner.

### Different lysine residues impact Cdc42p degradation in specific contexts

Ubiquitin modification involves the covalent modification of lysine residues by E3 ubiquitin ligases (Swatek and Komander, 2016). We previously identified lysine residues required for turnover of GTP-Cdc42p [Fig. 2A, K5, K94 and K96, red, (González and Cullen, IN PRESS)]. We next sought to identify lysine residues required for degradation of Cdc42p at 37°C. Systematic examination of lysine substitutions introduced into GFP-Cdc42p identified one lysine residue in the C-terminal region (K166) and four in the poly-basic C-terminal domain (PB) that were required for turnover of Cdc42p at 37°C (Fig. 2B, 37°C; *Fig. S2*). Other lysine residues were not required (Fig. 2C; levels, 37°C). In particular, the three lysine residues required for turnover of GTP-Cdc42p did not impact turnover of Cdc42p at 37°C (Fig. 2B, Q61L; TD, K5R, K94R, K96R). In line with this result, K166R and four lysines in the PB region were not required for degradation of GTP-Cdc42p (Fig. 2B, Q61L), indicating that each context requires non-overlapping lysine residues (Fig. 2, A and C). Some combinations of lysine substitutions did not promote the stability of the protein (Fig. 2C, K166R, K183R, K184R, K186R, K187R; K5, K123, K128, K166R; 12KR; 13KR), which indicates that lysines can play roles in protein stability and turnover (see below). Therefore, different lysine residues target Cdc42p for turnover in different settings.

**Figure 2.**
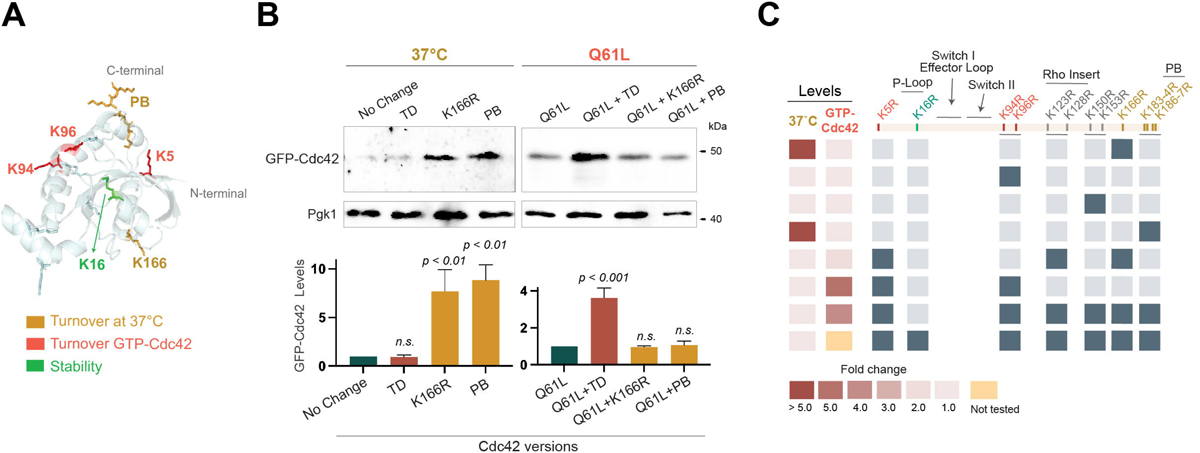
Lysines residues required for Cdc42p degradation at 37°C. **A)** The yeast Cdc42p protein sequence was overlaid onto the crystal structure of human Cdc42p. Yellow refers to lysines involved in Cdc42p turnover at 37°C, red refers to lysines involved GTP-Cdc42p (González and Cullen, IN PRESS) turnover and green indicates lysine involved in protein stability. **B)** 37°C, GFP-Cdc42p levels of wild-type cells (WT, ∑1278b, PC538) expressing GFP-Cdc42p (PC6454), GFP-Cdc42p^K5R,K94R,K96R^ (PC7636), GFP-Cdc42p^K166R^ (PC7518), GFP-Cdc42p^PB^ (PB: K183R, K184R, K186R, K187R; PC7520) grown and at 30°C for 5 h and shifted to 37°C for 2 h. Q61L, GFP-Cdc42p levels of wild-type cells expressing GFP-Cdc42p^Q61L^ (PC7458), GFP-Cdc42p^Q61L+TD^ (TD: K5R, K94R, K96R; PC7654), GFP-Cdc42p^Q61L,K166R^ (PC7638), GFP-Cdc42p^Q61L+PB^ (PC7665) grown and at 30°C for 5 h and shifted to 37°C for 2 h. See Fig. 1D for details. **C)** Heat map shows protein levels of indicated lysine substitutions of GFP-Cdc42p in wild-type cells. Color code represents the relative fold change of GFP-Cdc42p lysine substitutions compared to wild-type GFP-Cdc42p. Grey color indicates lysines substitutions to arginines in GFP-Cdc42p. Colors of alleles are described in Fig. 2A. Some data comes from González and Cullen IN PRESS.

### Cdc42p is turned over at 37°C by two pathways: by the proteosome and by the vacuole in an ESCRT-dependent manner

Proteins can be degraded by intracellular trafficking in the lysosome called the vacuole in yeast (Perera and Zoncu, 2016) and by the 26S proteasome (Grice and Nathan, 2016). Although Cdc42p is a cytosolic protein (which favors proteosome degradation), it also is modified by a lipid anchor and is associated with membranes (favoring vacuolar degradation). At 37°C, GFP-Cdc42p accumulated in the *cim3-1* mutant (also known as *rpt6-1*), which is defective for proteosome function (Ghislain et al., 1993) (Fig. 3A; similar data was reported in Gonzalez and Cullen, IN PRESS). *PEP4* encodes a vacuolar protease that is required for maturation and activation of vacuolar proteases (Parr et al., 2007). The *pep4*Δ mutant also showed elevated Cdc42p levels at 30°C and 37°C (*Fig. S3A*). Time course analysis revealed that degradation of GFP-Cdc42p levels at 37°C was slower in the *pep4*Δ mutant than wild type (Fig. 3B), indicating that the vacuole is required for Cdc42p degradation under physiological conditions and upon heat stress (37°C). Immunoblot analysis also showed that cleavage of GFP from GFP-Cdc42p, which is commonly mediated by vacuolar proteases (Hettema et al., 2004), was dependent on Pep4p (*Fig. S3B*). In line with this observation, the localization of GFP-Cdc42p was also impacted in cells lacking Pep4p. GFP-Cdc42p localized at the plasma membrane and internal membranes as previously reported (Etienne-Manneville, 2004). GFP-Cdc42p accumulated inside the vacuole in the *pep4*Δ mutant at 30°C (*Fig. S3C*). Taken together, these results indicate that Cdc42p can be turned over by the 26S proteasome and also by a mechanism that involves trafficking of the proteins for turnover in the vacuole.

**Figure 3.**
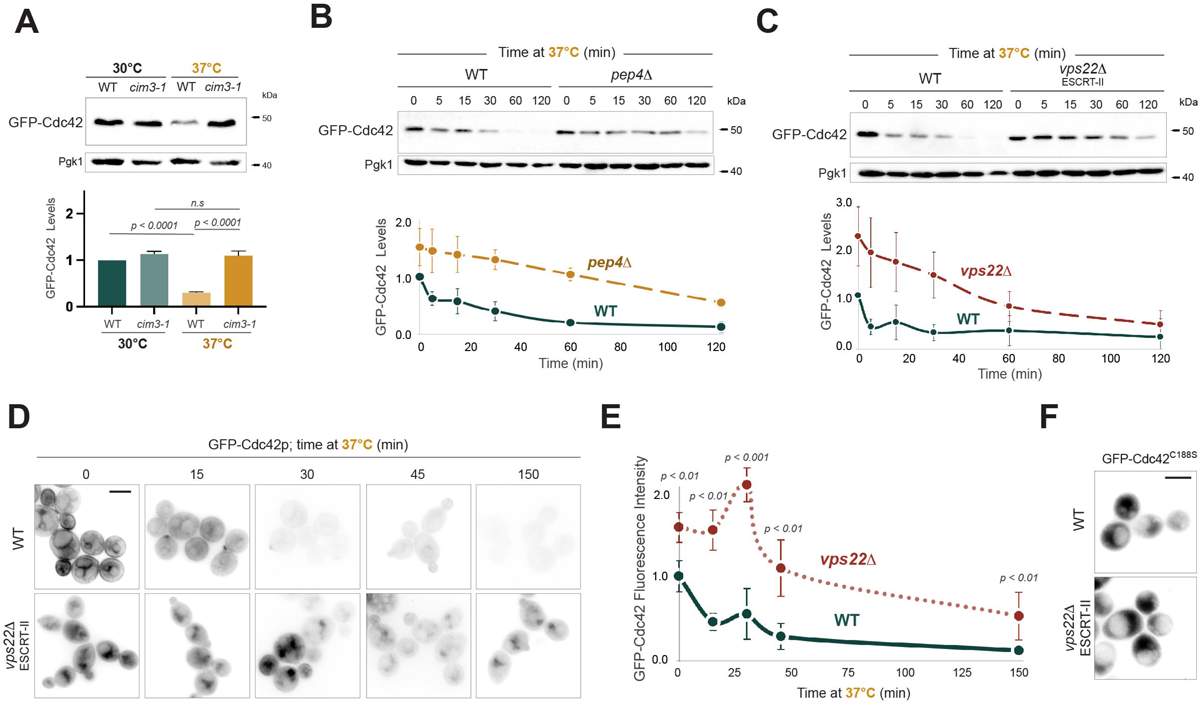
Cdc42p can be degraded by the vacuole in an ESCRT-dependent manner. **A)** Wild-type (WT, S288C, PC5851) cells and the *cim3-1* (PC5852) mutant expressing GFP-Cdc42p (PC6454) grown at 30°C for 5 h (30°C) and shifted to 37°C for 2 h (37°C). See Fig. 1D for details. **B)** Wild-type cells (WT, ∑1278b, PC999) and the *pep4*Δ mutant (PC3154) expressing GFP-Cdc42p (PC7354) grown at 37°C during the indicated time points. Bottom graph represents the quantification of GFP-Cdc42p levels compared to Pgk1p. Error bars represents S. D. from two biological replicates. **C)** Wild-type cells (WT, ∑1278b, PC6016) and the *vps22*Δ mutant (PC7548) expressing GFP-Cdc42p (PC6454) grown at 37°C during the indicated time points. See Fig. 3D for details. **D)** Wild-type cells and the *vps22*Δ mutant expressing GFP-Cdc42p (PC6454) were incubated and 30°C for 5h (30°C) and shifted to 37°C, cells were visualized at the indicated time points. Bar, 5µm. **E)** Relative fluorescence quantification of same cells. Data were analyzed by one-way ANOVA followed by Tukey’s multiple comparison test, n = 25, S.D. represents the standard deviation. **F)** Fluorescence microscopy of wild-type cells and the *vps22*Δ mutant expressing GFP-Cdc42p^C188S^ (PC7350) grown to mid-log phase at 30°C. Bar, 5µm.

We further examined the role of the trafficking pathway on Cdc42p degradation at 37°C. The ESCRT complex regulates the invagination of proteins from the surface of endosomes to the MVB (Remec Pavlin and Hurley, 2020). MVB fusion with the vacuole leads to the turnover of cargo proteins by vacuolar proteases (Henne et al., 2011; Schmidt and Teis, 2012). ESCRT is made up of several protein complexes [ESCRT-0, ESCRT-I, ESCRT-II, ESCRT-III and the Vps4p complex], and the structure and mechanism of action of ESCRT has recently been elucidated (Banjade et al., 2021; Bertin et al., 2020; Maity et al., 2019; Remec Pavlin and Hurley, 2020; Strohacker et al., 2021). We examined the levels of Cdc42p in cells lacking Vps22p which is a component of the ESCRT-II complex (Hierro et al., 2004). At 37°C, turnover of GFP-Cdc42p was defective in cells lacking Vps22p during the first 60 min (Fig. 3C; *Fig. S3D*), which indicates that ESCRT is required for Cdc42p degradation at initial time points. However, GFP-Cdc42p was degraded at 120 min in cells lacking Vps22p, suggesting that prolongated incubation of cells at 37°C induces the degradation of Cdc42p in an ESCRT-independent manner. Similar to Vps22p, prolonged incubation at 37°C induced Cdc42p degradation in the *pep4*Δ mutant (Fig. 3B; 120 min). These results indicate that the vacuole contributes to Cdc42p turnover at 37°C; however, the proteasome makes a more prominent contribution.

Time-lapse fluorescence microscopy also showed accumulation of GFP-Cdc42p in cells lacking Vps22p (Fig. 3D). Quantification of the fluorescence signal showed that the reduction in GFP-Cdc42p levels at 37°C was dependent on Vps22p (Fig. 3E). In addition, we also compared the turnover of Cdc42p in ESCRT mutants to the turnover of Msb2p-GFP, a transmembrane signaling glycoprotein that is known to be degraded in an ESCRT- and Pep4p-dependent pathway (Adhikari et al., 2015b) (*Fig. S3E*). We found that GFP-Cdc42p levels were slightly stabilized in several ESCRT mutants at 30°C (*Fig. S3F*). Therefore, the 26S proteasome and the trafficking pathway (an ESCRT-dependent pathway that culminates in the vacuole) regulate the degradation of Cdc42p under physiological and heat stress (37°C) conditions.

Rsp5p mediates the turnover of plasma membrane proteins through the trafficking pathway to the vacuole (Gajewska et al., 2001; Lauwers et al., 2010). Cdc42p is associated with the plasma membrane by prenylation, which occurs by modification of the cysteine at position C188 (Ziman et al., 1991). We tested whether a version of Cdc42p that cannot be prenylated, GFP-Cdc42p^C188S^, was capable of being turned over in the trafficking pathway. Cells lacking Vps22p did not influence GFP-Cdc42p^C188S^ localization (Fig. 3F), and the cleavage of GFP from GFP-Cdc42p^C188S^ was significantly reduced (*Fig. S3G*). These results indicate that Cdc42p localization at the plasma membrane is critical for its internalization to endosomes and for ESCRT-dependent turnover in the vacuole. Some ESCRT components function in a pathway that senses changes in pH, called the RIM101 pathway (Xu et al., 2004). The RIM101 pathway can regulate the Cdc42p-dependent MAPK pathway that controls filamentous growth (Chavel et al., 2014), and therefore it was important to test whether the RIM101 pathway was involved in regulating Cdc42p localization and turnover. GFP-Cdc42p localization was not impacted in cells lacking Rim8p (*Fig. S3H*), which excludes the RIM101 pathway as regulator of Cdc42p localization and turnover. Taken together, these data show that a Rho-type GTPase can be turned over in by an ESCRT-dependent vacuolar pathway.

### The ESCRT pathway and vacuole are not required for turnover of GTP-Cdc42p

We previously showed that GTP-Cdc42p is turned over by the proteasome (González and Cullen, IN PRESS). The fact that Cdc42p is also turned over in the trafficking pathway, prompted us to investigate whether the turnover of GTP-Cdc42p might also occur in the ESCRT-to-vacuole pathway. We compared localization of GFP-Cdc42p to Msb2p-GFP. In wild-type cells, as previously shown (Adhikari et al., 2015b), Msb2p-GFP was mainly found inside the vacuole due to the high rate of turnover of the protein (Fig. 4A, WT, Msb2p-GFP), colocalized with the FM4-64 dye in the MVB in cells lacking Vps22p (Fig. 4A, *vps22*Δ, Msb2-GFP) and accumulated in the MVB in cells lacking several ESCRT components (Fig. 3B, left, Msb2-GFP). GFP-Cdc42p colocalized with FM4-64 in wild-type cells and was trapped in the MVB in cells lacking Vps22p (Fig. 4A, *vps22*Δ, GFP-Cdc42). GFP-Cdc42p also piled up in MVB in several ESCRT components (Fig. 4B, middle, GFP-Cdc42), corroborating that ESCRT is required for Cdc42p localization to MVB. However, GFP-Cdc42p^Q61L^, which tightly localizes to the plasma membrane [Fig. 4BA WT, GFP-Cdc42^Q61L^; (Woods et al., 2016a)], did not show colocalization with FM4-64 in cells lacking Vps22p (Fig. 4C, *vps22*Δ, GFP-Cdc42^Q61L^) or in any ESCRT component examined (Fig. 4B, left, GFP-Cdc42^Q61L^). One component of ESCRT (Vps27p, ESCRT-0) was not required for Msb2p-GFP or GFP-Cdc42p degradation (*Fig. S3C-D*), and Msb2-GFP was found inside of the vacuole in this mutant (*Fig. S3I*), thus, we did not further examine GFP-Cdc42p localization in cells lacking ESCRT-0 components. Consistently, the level of a GTP-locked version of GFP-Cdc42p, GFP-Cdc42p^Q61L^, was not stabilized in cells lacking Pep4p at either temperature (Fig. 4C, compare to *Fig. S3A*). In fact, the *pep4Δ* mutant showed reduced levels of Cdc42p^Q61L^ compared to wild-type cells, which may occur because cells lacking Pep4p may be more efficient at degrading Cdc42p by the proteasome. By immunoblot analysis, the cleavage of GFP (which is characteristic of vacuolar degradation) from GFP-Cdc42p^Q61L^ was reduced compared to wild type (*Fig. S3J*; Q61L). Taken together, these results indicate that the active conformation of Cdc42p, GTP-Cdc42p, is not turned over by ESCRT in the trafficking pathway.

**Figure 4.**
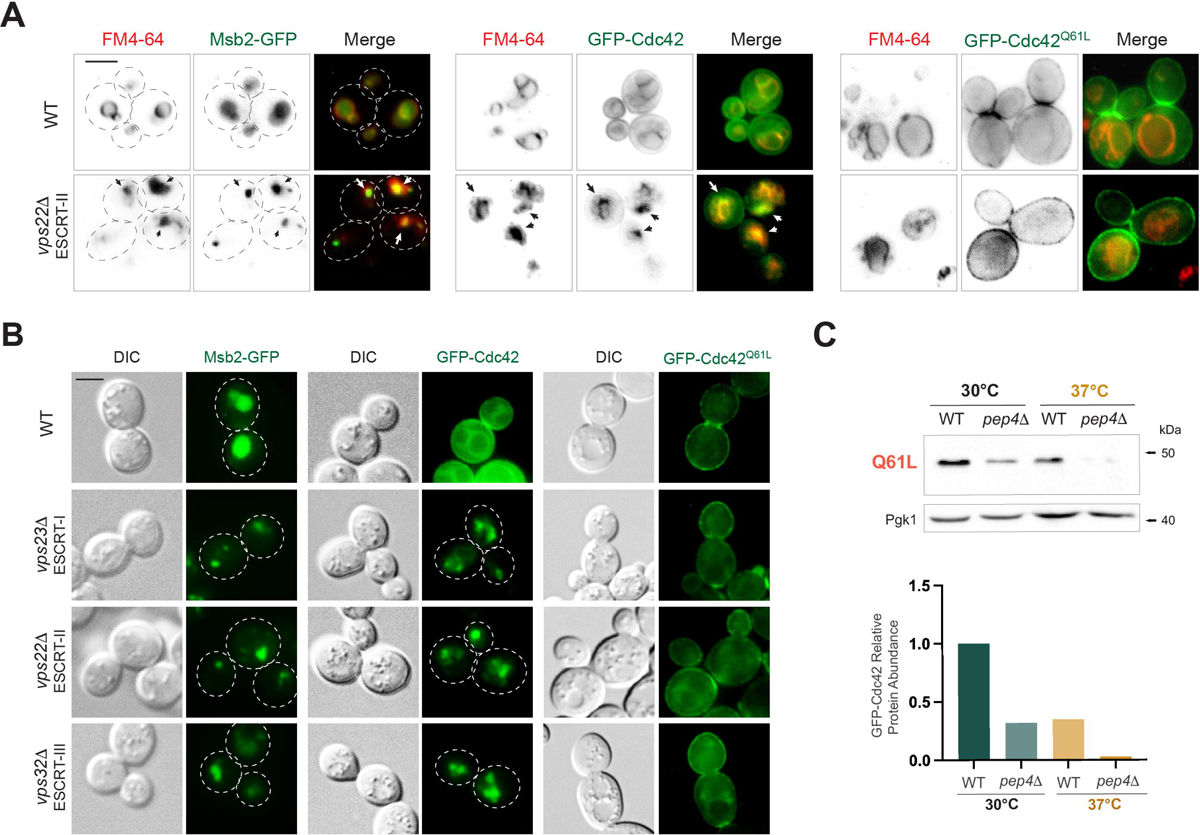
GTP-Cdc42p is not degraded by the ESCRT-vacuole pathway. **A)** Fluorescence microscopy of wild-type cells (WT, ∑1278b, PC6016) expressing Msb2p-GFP (PC2582), GFP-Cdc42p (PC6454) or GFP-Cdc42p^Q61L^ (PC7458) grown for 5 h at 30°C and stained with FM4-64 dye. Arrows indicate colocalization of GFP-Cdc42p and FM4-64. Bar, 5µm. **B)** Fluorescence microscopy of wild-type cells (PC6016) and the indicated ESCRT mutants expressing Msb2p-GFP, GFP-Cdc42p, or GFP-Cdc42p^Q61L^ grown for 5 h at 30°C. Bar, 5µm. **C)** Wild-type cells (WT, S288C, PC986) and the *pep4*Δ (PC3063) mutant expressing GFP-Cdc42p^Q61L^ grown at 30°C for 5 h (30°C) and shifted to 37°C for 2 h (37°C). See Fig. 1D for details.

### Turnover of Cdc42 at 37°C impairs the response to mating pheromone and impacts cell growth

High temperatures might induce Cdc42p mis-folding, resulting in turnover of the inactive version of the protein. Alternatively, Cdc42p turnover at high temperatures might be a regulated response that influences one or more of its functions in the cell. To test this possibility, we first looked at the mating response, which is regulated by a Cdc42p-dependent MAPK pathway (Bardwell, 2005; Chen et al., 2000; Chou et al., 2004; Good et al., 2009; Herskowitz, 1995; Kim and Rose, 2022; Li et al., 2017; Metodiev et al., 2002; Velazhahan et al., 2021). The addition of α-factor causes cell cycle arrest in the G1 phase of the cell cycle (halo formation) and can be used to measure the activity of the mating pathway (Butty et al., 1998; Long et al., 1997; Nern and Arkowitz, 1999). We found that halo size declined as temperature increased (Fig. 5, A and B), indicating that mating is compromised at high temperatures. This could be a result of Cdc42p turnover at high temperatures. Pheromone sensitivity was restored in cells lacking Ydj1p (Fig. 5, A and B), which have elevated levels of Cdc42p (Fig. 1D). Taken together these results suggest that Cdc42p turnover at high temperature impacts Cdc42p functions.

**Figure 5.**
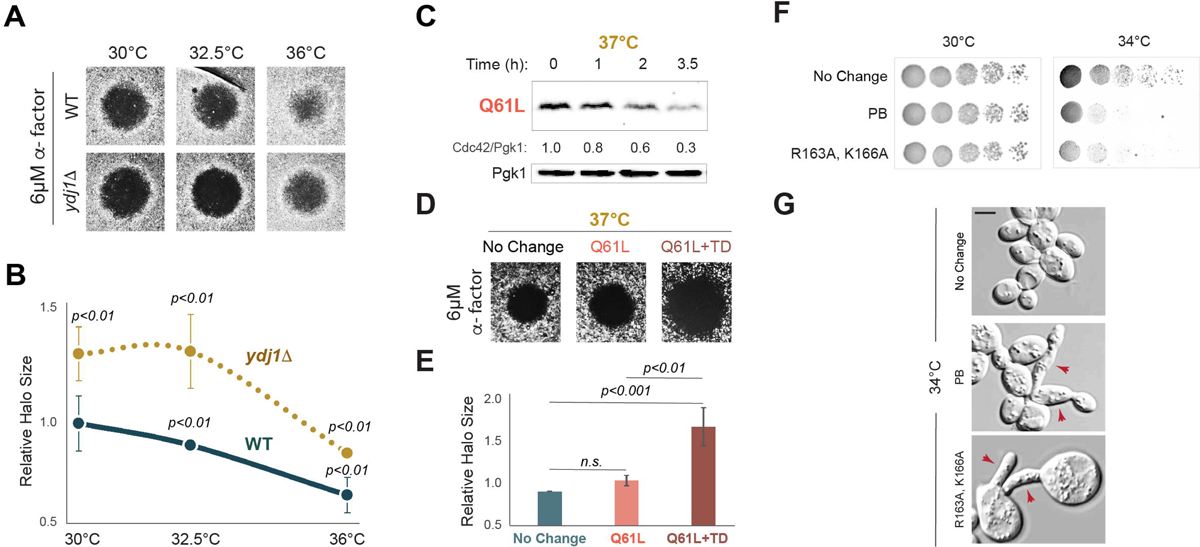
Defective Cdc42p turnover at 30°C impact MAPK activity and cell polarity. **A)** Halo formation in response to α-factor of wild-type (WT, S288C, PC896) cells and the *ydj1*Δ (PC7657) mutant. Cells grown for 16 h were spread on SD media where 6 µM α-factor was spotted on the top to study cell cycle arrest, plates were incubated for 48 h at the indicated temperature. **B)** Relative halo size of same cells examined in panel 3A. Data were analyzed by one-way ANOVA followed by a Tukey’s multiple comparison test, n = 4, Error bars refer to S.D. **C)** Levels of GFP-Cdc42p^Q61L^ in WT cells incubated at 37°C for the indicates times. Cdc42/Pgk1 ratio refers to relative levels of GFP-Cdc42p^Q61L^ to Pgk1p. **D)** Halo formation in WT cells expressing GFP-Cdc42p (Cdc42), GFP-Cdc42p^Q61L^ (Q61L), and GFP-Cdc42p^K5R; Q61L; K94R; K96R^ (Q61L+TD) in response to two concentrations of a-factor, 1.6 µM and 6 µM. Cells were incubated for 48 h at 37°C. **E)** Bottom, data were analyzed by one-way ANOVA followed by a Tukey’s multiple comparison test, n= 4, Error bars refer to standard deviation. **F)** Serial dilutions of wild-type cells and *CDC42* alleles grown at 30°C and 34°C for 3 days. **G)** Wild-type cells, Cdc42p^K183A,K184A,K186A,K187A^ (PB), and Cdc42p^K163R,K166R^ alleles were grown on YEPD for 48 h at 34°C. Arrows indicate hyper-polarized cells. Bar, 5µm.

We next asked whether Cdc42p stabilization by lysine substitutions impacts the mating response. Lysine substitutions in the poly-basic C-terminus (PB) did not significantly impact halo formation, but K166A (also containing R163A) caused the formation of larger haloes specifically at high temperatures (*Fig. S4, A-B*; 34°C), which corroborates the idea that Cdc42p might be degraded at high temperatures to impact its functions in the cell. In line with this possibility GTP-locked Cdc42p is also turned over at 37°C, albeit with a different kinetic profile (Fig. 5C). GTP-Cdc42p does not impact mating at 30°C [González and Cullen, IN PRESS, *Fig. S4C*, 30°C)]. However, a turnover-defective (TD) version of GTP-locked Cdc42p, Cdc42p^Q61L+TD^ (Cdc42p^Q61L,K5R,K94R,K96R^), also formed larger haloes at 37°C (Fig. 5, D and E). This result reinforces the idea that turnover of GTP-Cdc42p attenuates mating at 37°C. It should be noted that the K5R, K94R, and K96R did not impact total Cdc42p levels at 37°C (Fig. 2B), suggesting that these three lysines might be exclusively targeted for degradation of the GTP-bound conformation, either at 30°C or 37°C. Cdc42p also regulates the fMAPK and HOG pathways, which were not tested here.

Cdc42p stimulates bud growth through the binding to different effector proteins, which is coordinated with the cell cycle (Miller et al., 2020; Moran et al., 2019). We next explored whether preventing Cdc42p degradation at high temperatures impacts cell viability. Cdc42p^PB^ and Cdc42p^R163A,K166R^ mutants were defective for growth at high temperatures (Fig. 5F; 34°C). Cells defective for Cdc42p turnover at 34°C were examined by microscopy. Interestingly, cells defective for Cdc42p turnover at 37°C showed morphological defects compared to wild-type Cdc42p (Fig. 5G; PB; R163A, K166A; more examples in *Fig. S4D*). These included increased cell size and hyper-polarized growth (Fig. 5G; *Fig. S4D*), which are different than phenotypes described for accumulation of GTP-Cdc42p (Gonzalez and Cullen IN PRESS). Taken together, these results indicate that turnover of Cdc42p at high temperatures has functional consequences in a subset of Cdc42p-dependent processes. Failure to degrade Cdc42p in this setting has detrimental consequences on cell viability and cell polarity.

### Lysine residue K16 is required for stability

By analyzing the levels of versions of Cdc42p containing lysine substitutions, we found that some versions showed reduced levels compared to wild type (Fig. 6A; *Fig. S5A*). These included eight versions of the protein tested that contained lysine substitutions in different parts of the protein (Fig. 6B). The low levels of Cdc42p presumably result from enhanced turnover. We traced the stability defect of Cdc42p to a single residue, K16, a lysine residue that is located in the P-loop of the protein and which is not surface exposed. Versions of Cdc42p that contained the substitution K16R were found at low levels in the cell (Fig. 6, A-C, shown in green).

**Figure 6.**
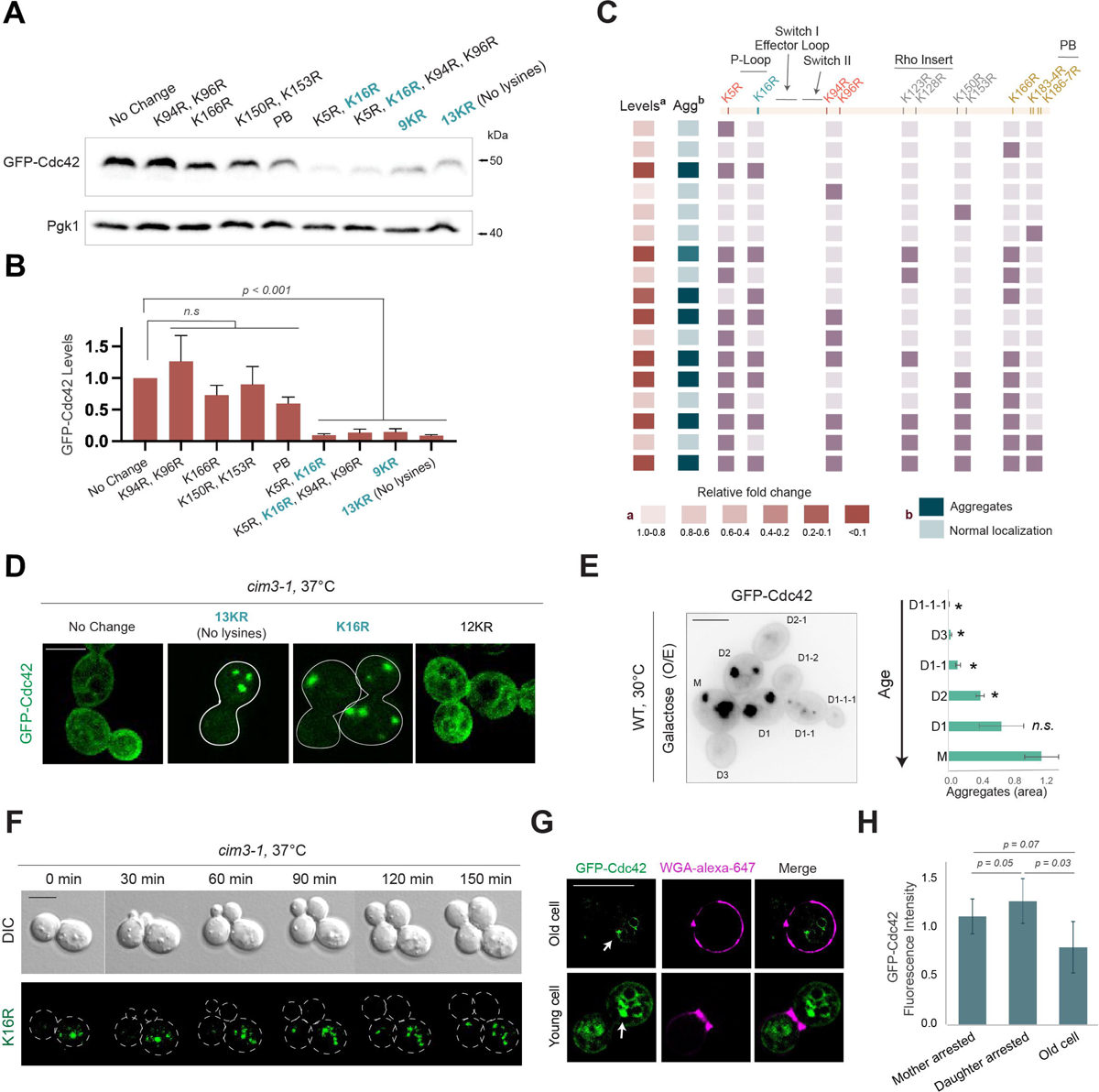
Analysis of the K16R mutation: impact on Cdc42p levels and aggregate formation. Cdc42p aggregates mostly localized in aging cells. **A)** Wild-type cells (WT, ∑1278b, PC538) expressing GFP-Cdc42p (No change, PC6454), GFP-Cdc42p^K94R,K96R^ (PC7508), GFP-Cdc42p^K166R^ (PC7518), GFP-Cdc42p^K150R,K153R^ (PC7511), GFP-Cdc42p^K183R,K184R,K186R,K187R^ (PC7520), GFP-Cdc42p^K5R,K16R^ (PC7504), GFP-Cdc42p ^K5R,K16R,K94R,K96R^ (PC7507), GFP-Cdc42p^K5R,K16R,K94R,K96R,K123R,K128R,K153R,K166R^ (PC7513), and GFP-Cdc42p^K5R,K16R,K94R,K96R,K123R,K128R,K150R,K153R;,K166R,K183R,K184R,K186R,K187R^ (13KR; PC7521) grown to mid-log phase at 30°C. B) Quantification of GFP-Cdc42p levels from three biological replicates, n= 3, analyzed by one-way ANOVA and Tukey’s multiple comparison test. Error bars refers to standard deviation. C) Heat map showing aggregates resembling pattern of the indicated alleles expressed in *cim3-1* cells (Agg) and relative protein levels of indicated alleles expressed in wild-type cells grown to mid-log phase at 30°C (Levels). D) Confocal microscopy of *cim3-1* cells expressing GFP-Cdc42p (No change), GFP-Cdc42^K5R,K16R,K94R,K96R,K123R,K128R,K150R,K153R,K166R,K183R,K184R,K186R,K187R^ (13KR), GFP-Cdc42pK16R (PC7607), and GFP-Cdc42^K5R,K94R,K96R,K123R,K128R,K150R,K153R,K166R,K183R,K184R,K186R,K187R^ (12KR; PC7633) grown at 37°C for 2 h. Bar, 5µm. E) Left, fluorescence microscopy of wild-type cells expressing pP_GAL1_-GFP-linker-*CDC42P* grown for 8 h in YEP-GAL media. Bar, 5 µm. Right, relative quantification of the aggregates area from cells in Fig. 6E and *S6D*. F) Time-lapse confocal microscopy of the *cim3-1* mutant (PC5852) expressing GFP-Cdc42p^K16R^ (K16R) and grown at 37°C. Bar, 5µm. G) Fluorescence microscopy of MEP cells (PC7713) expressing GFP-Cdc42p (PC6454) and stained with WGA-alexa-647 grown for 48h at 30°C. Arrows indicate GFP-Cdc42p aggregates. Bar, 5µm. H) Relative quantification of GFP-Cdc42p levels at the plasma membrane of MEP cells grown for 48h at 30°C. p-values were obtained using Mann-Whitney U test, n > 25 cells.

Incidentally, the fact that GFP-Cdc42p^13KR^ was stabilized in the in the *cim3-1* proteasome mutant (*Fig. S5B*) was intriguing because Cdc42p^13KR^ does not contain any lysine residues. This version of the protein was also present at low levels in the cell (Fig. 6B, 13KR). This data suggests that this version of Cdc42p can be turned over in a manner that does not require lysine residues, which is unexpected because protein ubiquitination typically occurs on lysine residues (Hochstrasser, 2000). To confirm this possibility, the GFP epitope, (which does contain lysine residues), was substituted by a Hisx6 tag. The Hisx6-Cdc42p^13KR^ protein that lacks any lysine residues was also present at low levels (*Fig. S5C*), indicating that it is turned over in a lysine-independent manner. In some proteins, cysteine, serine, and threonine can act as ubiquitin acceptors (Kravtsova-Ivantsiv and Ciechanover, 2012; Lei et al., 2018; McClellan et al., 2019; McDowell and Philpott, 2013; Tait et al., 2007; Wang et al., 2007).

### Accumulation of Cdc42p^K16R^, or overexpression of the protein, leads to the formation of aggregates

The localization of unstable versions of GFP-Cdc42p was examined by fluorescence microscopy, which confirmed that these proteins are present at low levels in the cell (Fig. 6B). The localization of these proteins was also examined in the *cim3-1* mutant at 37°C, where these unstable proteins would be expected to accumulate. Compared to wild-type GFP-Cdc42p (Fig. 6A; GFP-Cdc42 ≥ 0.5), which showed a normal localization pattern (Fig. 6C), versions of Cdc42p that were found at reduced levels (Fig. 6D; *Fig. S5D*; GFP-Cdc42 ≤ 0.2) showed a punctuate pattern indicative of protein aggregates (Fig. 6C). For example, Cdc42p^13KR^ accumulated in cells lacking a functional proteasome based on immunoblot analysis (*Fig. S5B*) and formed aggregates in the *cim3-1* mutant (13KR; Fig. 6, C and D). Turnover of Cdc42p^13KR^ was also dependent on Ydj1p (*Fig. S5E*). Therefore, the turnover of Cdc42p^13KR^ occurs in the 26S proteasome to inhibit the accumulation of protein aggregates.

As for protein stability, the formation of aggregates of GFP-Cdc42p^13KR^ was traced to the lysine residue, K16. Cdc42p^K12R^, which lacks all of the lysines in Cdc42p except K16, showed a normal localization pattern (Fig. 6D; 12KR). Likewise, GFP-Cdc42p^K16R^, which contains only a single lysine substitution at position K16, formed aggregates (Fig. 6D; K16R). We furthermore saw a direct correspondence between versions of Cdc42p containing K16R, reduced protein levels (< 0.2), and aggregate formation (Fig. 6C). Given there are few examples of aggregate formation of Rho-type GTPases, this phenotype was examined in more detail.

We first asked whether wild-type versions of Cdc42p were capable of forming aggregates. Growth at 37°C did not induce aggregation of GFP-Cdc42p (Fig. 6D, no Change; *Fig. S5D*) but overexpression of the *GFP-CDC42* gene from a strong inducible promoter (P*_GAL1_*) induced a punctate localization pattern in some cells (Fig. 6E, ∼20% of cells, t = 6h). As for K16R, aggregates induced by Cdc42p overproduction varied in number and size and showed a cytosolic pattern that was distinct from the lipophilic dye FM4-64 (*Fig. S6A*). Cells lacking Ydj1p also accumulated higher levels of the Cdc42p protein when *CDC42* was overexpressed (*Fig. S6B*) and aggregates were detected at earlier time points than seen in wild-type cells (*Fig. S6C,* t= 4h). For many proteins, aggregates are retained in mother cells, which occurs by a Ydj1p-(HSP40-) dependent mechanism (Saarikangas et al., 2017), presumably to ensure that properly folded proteins are enriched in daughter cells to promote cellular rejuvenation (Hill et al., 2017). When overexpressed, Cdc42p aggregates preferentially accumulated in mother cells (Fig. 6E; *Fig. S6D*). The filamentous strain background (∑1278b) was used to facilitate the assessment of cell lineage, because adhesive cells fail to separate. GFP-Cdc42p^K16R^ aggregates were also enriched in mother cells in the *cim3-1* mutant (Fig. 6F, see control in *Fig. S6E*; Movie 1, control GFP-Cdc42p; Movie 2, GFP-Cdc42p^K16R^).

We next explored the localization of GFP-Cdc42p in mother cells using the <u>M</u>other <u>E</u>nrichment <u>P</u>rogram (MEP) (Lindstrom and Gottschling, 2009; Moreno et al., 2019). We observed some GFP-Cdc42p aggregates in older cells, however, unexpectedly, GFP-Cdc42p also formed aggregates in young cells (Fig. 6G, arrows). Therefore, we could not establish a correlation between age and Cdc42p aggregates using this approach. Young cells in the MEP program lack the anaphase-promoting complex (APC) activator *CDC20*, which fails to degrade mitotic cyclins required for cell cycle progression (Visintin et al., 1997). Thus, aggregated Cdc42p in those cells might occur due to defective protein degradation of Cdc42p or perhaps due to general proteostatic stress. Aggregates in arrested cells did not colocalize with the disaggregate Hsp104p (*Fig. S6F*). Analysis of GFP-Cdc42p in older cells, also revealed low levels of GFP-Cdc42p at the plasma membrane compared to arrested daughter cells (Fig. 6H), which might be expected as these cells lose their ability to form buds. Although the formation of aggregates might in principle cause deleterious phenotypes, overexpression of GFP-Cdc42p did not cause a growth defect (*Fig. S6F*). Likewise, aggregates induced by K16R did not cause a noticeable growth defect (Movie 2). Therefore, Cdc42p can form aggregates when overproduced or when versions of the protein that compromise stability accumulate to high levels in the cell.

## DISCUSSION

Rho GTPases govern many aspects of cellular function including cytoskeletal dynamics and signal transduction pathways to control a wide range of biological responses. Altered Rho GTPase regulation impacts human health, including for example the induction of malignant transformation in some cancers (Goka and Lippman, 2015; Huang et al., 2013; Tamehiro et al., 2015; Zhang et al., 2019) and is an underlying cause of some neurological (Aguilar et al., 2017; Guiler et al., 2021) and immunological diseases (El Masri and Delon, 2021). Understanding the regulatory determinants of Rho GTPase stability and turnover can provide insights into how these proteins are regulated and may impact our understanding of the roles these regulatory features play in the healthy and disease states. By exploring the turnover regulation of the Rho GTPase Cdc42p in yeast, we have uncovered new and potentially general insights into aspects of Rho GTPase regulation that may extend to other systems.

### Rho GTPase turnover at high temperatures: identification of lysines and elucidation of the turnover mechanism

We previously showed that chaperones of the HSP40 and HSP70 family and the NEDD4-type ubiquitin ligase Rsp5p were required for turnover of GTP-Cdc42p, and we identified lysines involved in its turnover [TD: K5, K94, and K96 (González and Cullen, IN PRESS)]. Here, we show that Cdc42p is degraded in response to growth at high temperatures, which requires the same HSP40, HSP70, and Rsp5p proteins and a different set of lysine residues (K166; PB Fig. 7). The fact that GTP-Cdc42p requires different lysine residues than degradation of the protein at 37°C might be important, since GTP-Cdc42p degradation only impacts the active state of the protein (presumably a small fraction of the protein), while at 37°C most of the protein in the cell is degraded. Different lysines may target the protein for turnover perhaps because the conformation of Cdc42p might change in different ways in response to GTP binding and protein folding at high temperatures, which may impact lysine accessibility by Rsp5p, or recognition of different parts of the protein by HSPs. Alternatively, the exposure of lysines might be influenced by different Cdc42p-interacting proteins that bind Cdc42p in the two contexts. Cdc42p turnover at 37°C was mediated by the 26S proteasome and the vacuole. Given that that proteasomes become more active upon heat shock in mammalian cells (Lee and Goldberg, 2022), this temperature-dependent activation of proteosome function might favor the degradation of Cdc42p by the proteasome under this condition.

**Figure 7.**
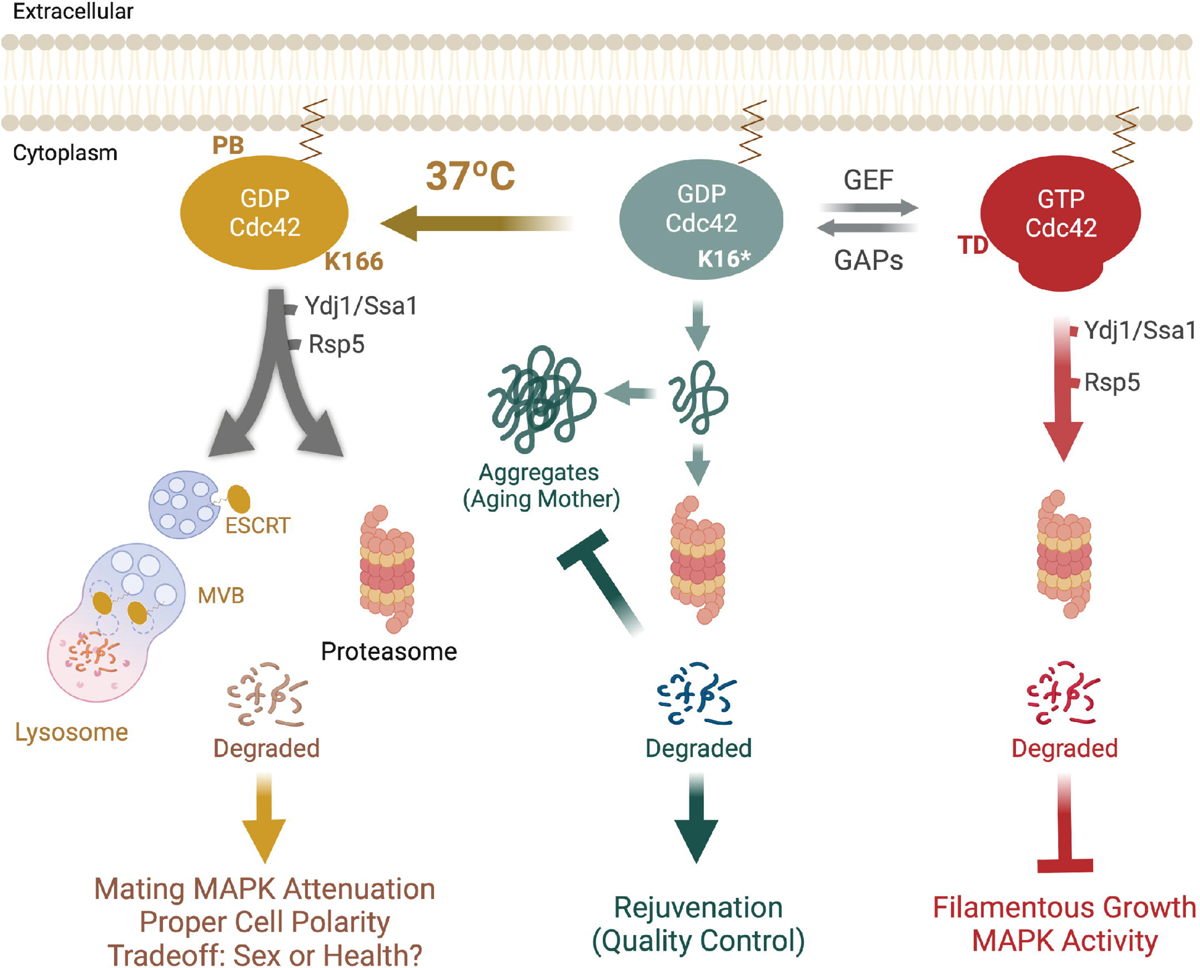
Cdc42p turnover in yeast. At 37°C, Cdc42p is degraded by the chaperones Ydj1p and Ssa1p (HSP40 and HSP70) and the NEDD4 ubiquitin ligase Rsp5p. Cdc42p turnover at 37°C requires K166, and lysines located at C-terminus of the protein (PB) and occurs in the proteasome and in the ESCRT-to-vacuole (lysosome) pathway. Cdc42p degradation at 37°C promotes cell health and inhibits the mating MAPK pathway. The K16 residue is critical for Cdc42p stability, mutation by K causes the protein to be degraded by the proteasome to inhibit formation of protein aggregates. Aggregates are confined in mother cells. As described in González and Cullen, IN PRESS, active Cdc42p is degraded by Ydj1p, Ssa1p and Rsp5 in the proteasome to attenuate the filamentous growth MAPK pathway.

Cdc42p is a highly conserved protein from yeast to humans. We have identified conserved lysines in the Cdc42p protein that are important for turnover of the protein at 37°C. Cdc42p^K166R^ is resistant to degradation at high temperatures. Based on cBioPortal for Cancer Genomics, K166 is a hotspot for cancer mutation in CDC42 (Crosas-Molist et al., 2022). It would be interesting to explore whether mutations at that site lead to elevated protein levels in malignant cells. We also identified residues in the polybasic domain that contribute to turnover. In humans, CDC42 has two splice variants (Hart et al., 1991; Olenik et al., 1999) that differ in nine amino acids at the C-terminus of the protein. CDC42 plays an essential role during early mammalian development (Chen et al., 2000), and recently it has been shown that the two isoforms perform antagonist functions during neural differentiation, in particular by controlling mTOR activity (Endo et al., 2020). It is not clear how almost identical proteins are specificality regulated to perform different functions in the cell. The lysines K183 and K184, which we found were required for degradation of Cdc42p at 37°C, are present in one splice variant but not another. K183 and K184 may target one variant of CDC42 for degradation, allowing the other variant to perform its functions in neural development. Generally speaking, mutations that lead to the upregulation of Rho GTPases might contribute to malignant transformation and cancer metastasis (Clayton and Ridley, 2020; Hodge et al., 2020) and potentially problems in development.

### The biological relevance of Rho GTPase turnover at high temperatures

Heat shock stress induce protein misfolding, and cells respond by promoting protein refolding and clearing damaged proteins by degradation in an HSP-dependent manner, since accumulation of misfolded protein can cause damage to cell function and viability (Klaips et al., 2018; Sala et al., 2017). Cdc42p may be turned over at high temperatures as a type of protein quality control mechanism to prevent protein misfolding. However, evidence here suggests that the turnover of Cdc42p at elevated temperatures also has functional relevance to biological processes. We specifically found that cells respond poorly to pheromone at elevated temperatures. Thus, cells may experience a tradeoff, favoring protein folding/stability at the expense of mating, which itself is a key driver of evolutionary fecundity. Turnover of active Cdc42p did not impact mating at 30°C, maybe because there are enough levels of the protein to reach the maximum activity required. However, at 37°C when the levels of Cdc42p are reduced, mating sensitivity also becomes reduced. Maybe this is a priority for Cdc42p functions, which depend on the available levels of Cdc42p. Interestingly, incubation at 37°C (body temperate) decreases a switch in cell type (from white to opaque cells) of the human pathogen *Candida albicans* that is required for mating (Johnson, 2003), indicating that mating deficient at 37°C might be a feature conserved across species.

Cdc42p turnover at 37°C might occur by recognition of the misfolded protein by HSP protein chaperones. However, preventing the degradation of Cdc42p at 37°C by deletion of Ydj1p or stabilization of Cdc42p by K166 mutation induced better activation of the mating pathway, which is regulated by Cdc42p. This indicates that at least some fraction of Cdc42p may be turned over to shut down the function of the protein. Preventing Cdc42p degradation at high temperatures negatively influenced cell viability. This suggests that degradation of Cdc42p at 37°C might be a coordinated response to inhibit any Cdc42p functions. At 37°C, the PKC pathway gets activated to repair cell wall damage in yeast (Levin, 2011). The Rho GTPase Rho1p, which shares downstream effectors with Cdc42p, regulates the PKC pathway to control aspects of polarized growth and morphogenesis (Andrews and Stark, 2000; Bettinger et al., 2007; Delley and Hall, 1999; Kono et al., 2012). During cytokinesis, Rho1p and Cdc42p function antagonistically (Atkins et al., 2013; Onishi et al., 2013; Tolliday et al., 2002). Thus, turnover of Cdc42p at 37°C may favor execution of a Rho1p-dependent response, to might ensure cell wall repair through the PKC pathway.

Intriguingly, the turnover of Cdc42p at 37°C may represent an example at the molecular level of the tradeoff between mating and cellular fitness. An ongoing goal of evolutionary biology is to identify the causes of sexual selection and diversification, including sexual selection, phenotypic diversity, and tradeoffs between mating and fitness in other contexts. Here, we unexpectedly found that cells sensitivity to mating pheromone is reduced at higher temperatures, suggesting that mating under this condition is not preferred. We show that was at least in part due to Cdc42p turnover under this condition. We further show that by restoring Cdc42p levels, we could restore pheromone sensitivities at higher temperatures, but this came at a cost to cell growth (or viability), corresponding to polarity defects. Therefore, Cdc42p activity promoting mating under one condition can lead to a trade off in overall fitness. This provides one of many such examples of tradeoffs to be identified (Johnston et al., 2013), here in this case shown by altering the levels of a regulatory GTPase in a condition-specific manner.

### An example of Rho GTPase turnover in the vacuole

Transmembrane proteins at the plasma membrane can be ubiquitinated for targeting by turnover in the secretory pathway. By comparison, cytosolic proteins are typically turned over by the proteosome. Rho GTPases represent a special case, because they are cytosolic proteins that associate with the plasma membrane by the addition of a lipid anchor. This feature may account for why Cdc42p is turned over by both the proteosome and in the trafficking pathways. Indeed, we found that versions of Cdc42p that cannot be lipid modified are not turned over by ESCRT. In both cases, the turnover of Cdc42p requires Rsp5p, which can function to control protein turnover in the cytoplasm, yet is also responsible for most if not all turnover of proteins at the plasma membrane.

Although this may be the first case for turnover of a Rho GTPase in the trafficking pathway, other types of GTPases follow a similar route. For example, RAS can also be degraded in a lysosome-(Lu et al., 2009) and proteasome-dependent manner (Jeong et al., 2018; Kim et al., 2009), which both lead to attenuation in MAPK signaling. The alpha subunit of heterotrimeric G-proteins, such as the mating pathway Gα subunit Gpa1p, also undergoes proteasomal and vacuolar degradation (Dohlman and Campbell, 2019). This differential regulation of Cdc42p might allow specificity in the degradation of Cdc42p in response to different stimuli. For example, GTP-Cdc42p is turned over mainly in the proteosome, whereas at 37°C, Cdc42p is turned over by both the proteosome and the vacuole. Alternative routes for Cdc42p degradation might be beneficial for accelerating Cdc42p turnover in specific contexts and in response to different stimuli. It will be interesting to determine whether the trafficking pathway is required for degradation of other Rho GTPases or CDC42 in humans and other systems.

### Determinants of Rho GTPase stability and the formation of protein aggregates in aging cells

Protein aggregates are assemblies formed by association of misfolded proteins mainly connected through hydrophobic interactions that accumulate in the intracellular environment (Balchin et al., 2016). Many cytosolic proteins and mRNAs can form aggregates, which can occur in a highly regulated manner in response to nutrient limitation and other stresses and which can also lead to cellular stress and aging (Alshareedah et al., 2020; Cabrera et al., 2020; Hill et al., 2017; Khong and Parker, 2020; Moreno et al., 2019; Samant et al., 2018). In mammals, accumulation of protein aggregates is a major cause of folding-related neurodegenerative disorders (Balchin et al., 2016; Dobson, 2002). We found that a single lysine residue in the Cdc42p protein that is not surface exposed (K16) was important for the stability of the protein. K16 has previously been shown to be required for Cdc42p function, as that mutation expressed endogenously is inviable (Kozminski et al., 2000). Thus, the instability of Cdc42p^K16R^ might account for its inability to function in the cell. When expressed in cells that fail to turn over Cdc42p (e.g. proteosome mutant), Cdc42^K16R^ forms protein aggregates. Despite the fact that many proteins have been shown to form cytosolic aggregates, few examples have been reported for Rho GTPases. Interestingly, high temperatures induce aggregation of another Rho GTPase, Rho1p^C17R^-GFP in fission yeast. In this case, the GFP tag inhibits the prenylation of Rho1p, and the protein forms aggregates when not targeted to the plasma membrane (Cabrera et al., 2020). Therefore, the stability of Rho GTPases, as well as their membrane localization, may contribute to the aggregation of the protein. We also found that overproduction of wild-type versions of Cdc42p can form aggregates. The formation of aggregates might be a mechanism to maintain Cdc42p protein levels within the stoichiometric limits forced by physical or functional protein interactions. Interestingly, Cdc42p aggregates – due to overexpression or misfolding (K16R) - were partitioned to mother cells, presumably as part of the protein rejuvenation response (Hill et al., 2017). Therefore, as is true for many protein quality control drivers (Santra et al., 2019), Rho aggregation may impact aging in cells. Furthermore, the same point mutation that causes Cdc42p aggregates, K16R, was found in patients with pancreatic cancer (cBioPortal for Cancer Genomics), which might suggest that alterations in that part of the protein are critical for its folding and stability.

## Supporting information

Figure S1. GFP-Cdc42p accumulated high molecular weight products in a Rsp5-dependent manner

Figure S3. Role of ESCRT-vacuole pathway in degrading Cdc42p

Figure S5. Cdc42p aggregate formation of lysine mutants.

Figure S6. Overexpression of Cdc42p caused aggregate formation.

## ABBREVIATIONS

CHX: cycloheximide
DIC: differential interference contrast
Glu: glucose; Gal, galactose
GAP: GTPase activating protein
GEF: guanine nucleotide exchange factor
GFP: green fluorescent protein
GTPase: guanine nucleotide triphosphate
HMW: high molecular weight
HR: homologous recombination
HSP: heat shock protein
kDa: kilodalton
MAPK: mitogen-activated protein kinase
MVB: multivesicular bodies
fMAPK: filamentous growth MAP kinase pathway
O.D.: PAK, p21-activated kinase
PB: polybasic
Rho: Ras homology
S. D.: standard deviation
SDM: site-directed mutagenesis
SDS-PAGE: sodium dodecyl sulfate-polyacrylamide gel electrophoresis
YEPD: yeast extract, peptone and dextrose
WT: wild type

## ACKNOWLEDGEMENTS

Thanks to Charles Boone (University of Toronto), Keith Kozminski (University of Virginia, Charlottesville, VA), Daniel Lew (Duke University, Raleigh NC), David Pellman (Harvard Medical School, Boston, MA), and Scott Emr (Cornell University, Ithaca, NY) for providing reagents. Thanks to Emily Mehle for assistance with experiments and lab members for suggestions. The work was supported by a grant from the NIH (GM098629).

## Author Contributions

B.G. designed experiments, generated data and wrote the paper. M.A. designed experiments. P.J.C. designed experiments and wrote the paper.

## MATERIALS AND METHODS

### Yeast strains, reagents and media

All Strains used in this study are listed in Table 1, plasmids in Table 2, and primers in Table 3. Yeast strains were grown in YEPD media [<u>Y</u>east <u>E</u>xtract <u>P</u>eptone <u>D</u>extrose; 1% bacto-yeast extract, 2% bacto-peptone, 2% dextrose], YEPGAL media (<u>Y</u>east <u>E</u>xtract <u>P</u>eptone <u>D</u>extrose; 1% bacto-yeast extract, 2% bacto-peptone, 2% galactose) media and SD media (Synthetic Dextrose; 0.67% yeast nitrogen base without amino acids, 2% dextrose) media supplemented with amino acids when required unless otherwise indicated (Rose et al., 1990). Cells were grown at 30°C or 37°C, otherwise temperature is indicated in the corresponding figure legend. To maintain the selection of the plasmids, cells were grown in SD media lacking uracil. pRS306-GFP-linker-Cdc42 (pDLB3609) plasmid was kindly provided by the Lew Lab (Woods et al., 2016b). To construct the pGFP-Cdc42 plasmid (PC6454) driven by *CDC42* promoter, pRS306-GFP-linker-CDC42 was subcloned with with *EcoR*I and *Sal*I into pRS316 [CEN/URA (Sikorski and Hieter, 1989)]. To generate the pHis6x-linker-Cdc42 and pHis6x-linker-Cdc42^13KR^ by homologous recombination, pGFP-Cdc42 was linearized with the restriction enzyme SnaBI (Cat#R0130S, New England Biolabs), and co-transformed in yeast with primers harboring the *HISX6* sequence and flanking regions to the *CDC42* promoter and linker, sequences listed in *Table 3.* Gene disruptions were done according to standard techniques using antibiotic resistance markers NAT, HYG and KanMX6 (Goldstein and McCusker, 1999; Longtine et al., 1998). GeneART^TM^ Site-Directed Mutagenesis (SDM) Kit (Cat#A13282, Thermo Fisher) was used to insert point mutations in the pRS306-GFP-linker-CDC42 plasmid. Other mutations were inserted in the pRS306-GFP-linker-CDC42 plasmid by *in vivo* homologous recombination in yeast. pRS306-GFP-linker-CDC42 plasmid was linearized with *PshA*I (Cat#R0593S, New England Biolabs) and co-transformed with a PCR product containing the desired K-to-R changes into an uracil auxotrophic strain (PC538). Positive isolates were confirmed by sequencing.

**Table 1.**
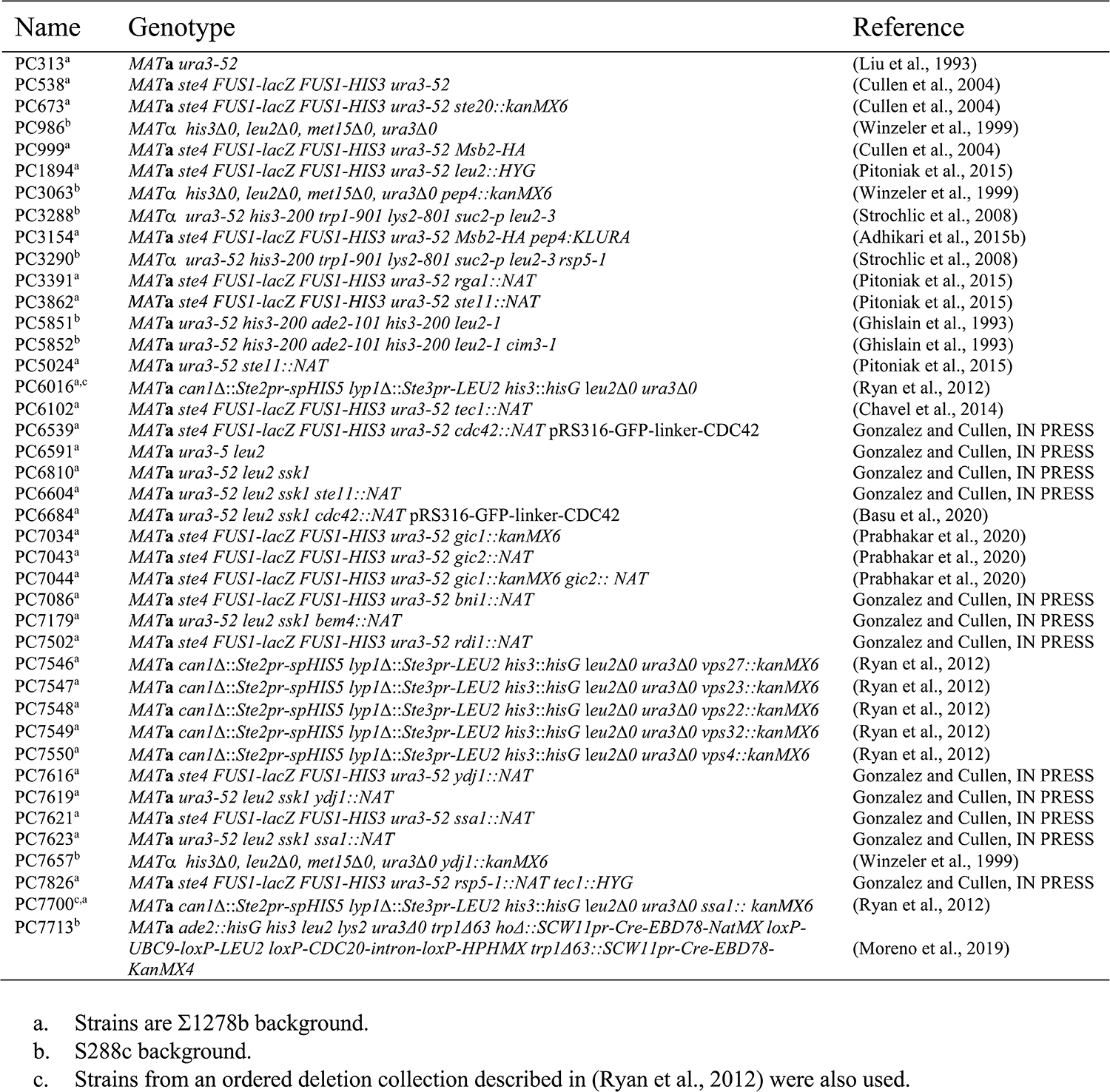
Yeast strains.

**Table 2.**
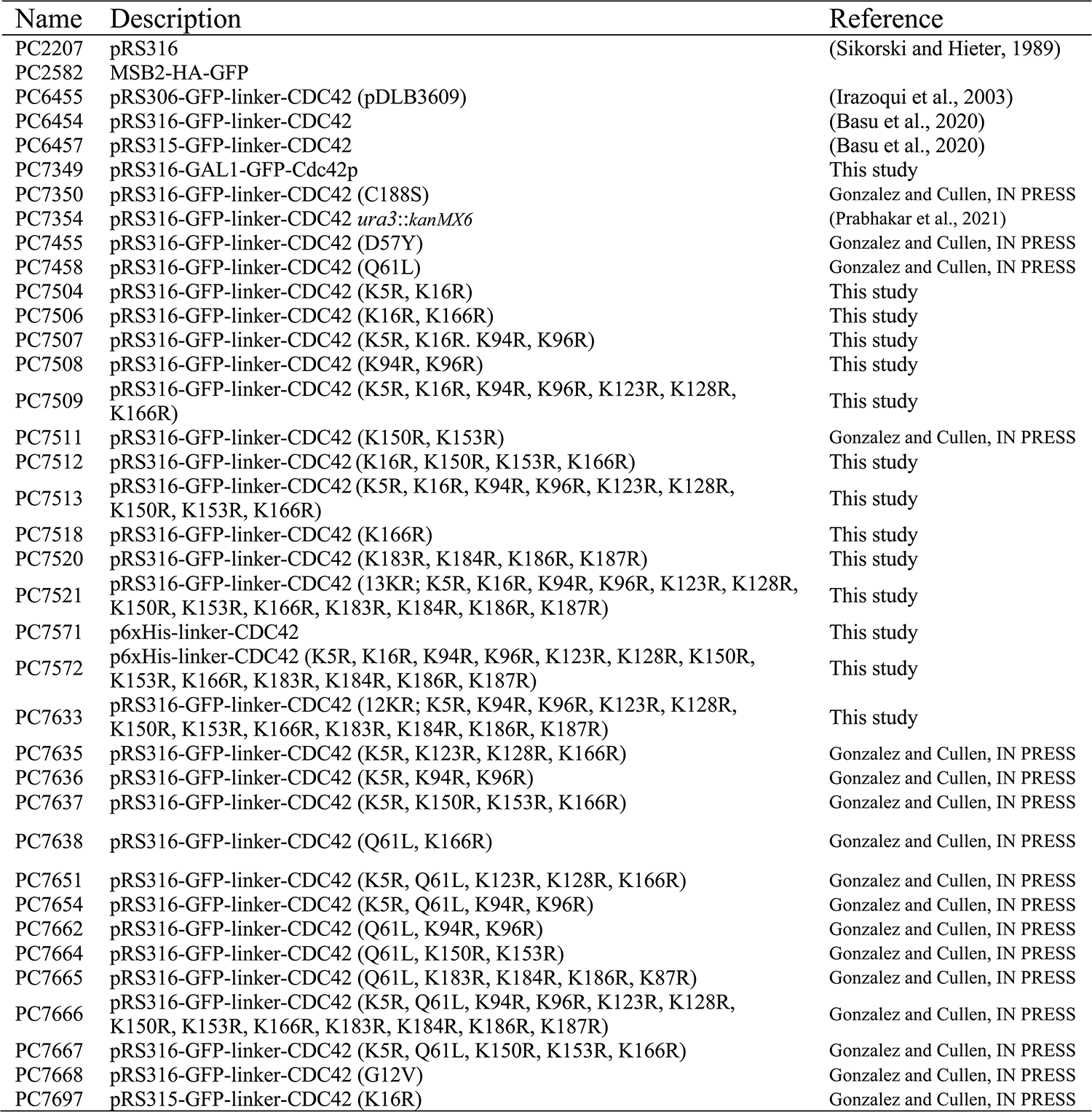
Plasmids used in the study.

**Table 3.**
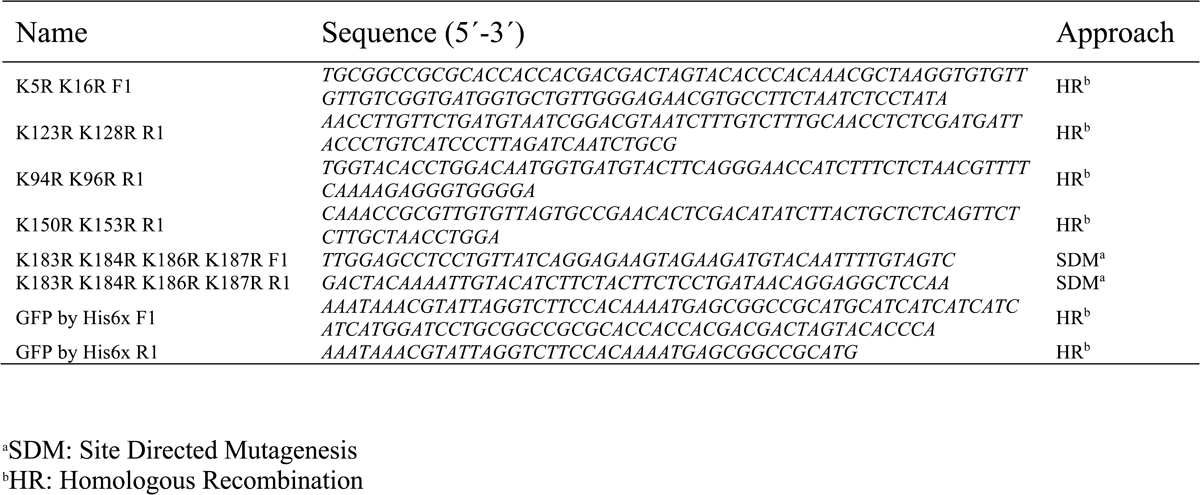
Primers used in the study.

### Protein immunoblot analysis

Cells were grown in YEPD or SD media for 16 h and resuspended into fresh media and grown for 4-6 h to mid-log phase or indicated time points. Immunoblotting was performed as described previously (Basu et al., 2020). In general, cell lysates were resolved by SDS-PAGE and proteins were transferred to a nitrocellulose membrane (Cat#10600003, Amersham^TM^ Protran^TM^ Premium 0.45 μm NC, GE Healthcare Life sciences) which were incubated in blocking buffer (5% nonfat dry milk, 10mM Tris-HCL [pH 8], 150mM NaCl, and 0.05% Tween 20) for 1 h at room temperature before antibody incubation. Incubations with the primary antibody were carried out at 4°C during 16h and the secondary antibody at room temperature for 1h. Antibodies were used at the manufacturer’s recommended concentrations. Proteins were visualized by Gel Doc XR Imaging System (Bio-Rad, Inc.), after addition of Chemiluminescent HRP substrate for chemiluminescent Westerns (Radiance^TM^ Plus Substrate, Azure Biosystems). Mouse monoclonal antibodies were used to detect green fluorescent protein (GFP) (Cat#11814460001, clones 7.1 and 13.1, Roche). Rabbit polyclonal anti-Cdc42p antibodies were provided by Dr. Keith Kozminski (Kozminski et al., 2000) and used at 1:1,000 dilution. Mouse monoclonal anti-Pgk1p antibodies (22C5D8, Cat#459250, Invitrogen) were used to measure protein levels. Anti-mouse IgG-HRP (Cat# 1706516, Bio-Rad Laboratories) and goat anti-rabbit IgG-HRP (Cat#115-035-003, Jackson ImmnunoResearch Laboratories) secondary antibodies were used. Quantification of band intensity were performed under non-saturated conditions and normalized to the Pgk1p protein using the Image Lab Software (Bio-Rad, Inc.).

Analysis of protein turnover with cycloheximide (CHX) was performed as previously described (Adhikari et al., 2015a). Briefly, 100ml of wild-type cells at 0.02 of O.D. were grown in SD media at 30°C. After 4h of incubation, the media was supplemented with 25 μg/ml of CHX, and 10 mL of samples were collected at 0, 15, 30, 45, 60, 90 and 120 min to generate cell extracts for immunoblot analysis. Experiments were performed in two biological replicates.

### Mating Pathway Activity and the Response to Pheromone

Mating pathway activity was evaluated by halo assays (Sprague et al., 1983). Cells were grown for 16 h in SD-URA media, and 200 μL of a 1:100 dilution were spread on SD-URA plates. After the plates were air dried, two concentrations of α-factor, 3 μL (1.8 μM) and 10 μL (6 μM) were applied to the surface. Plates were incubated at 30°C, 32.5°C, 34°C and 36°C for 2, 3 or 4 days and photographed.

### Modeling Protein Structure

To obtain the Cdc42p yeast structure, the protein sequence was overlapped onto the crystal structure of human Cdc42p using the Expasy web server SWISS-MODEL (https://swissmodel.expasy.org) (Nassar et al., 1998).

### cBioPortal for Cancer Genomics

The website of cBioPortal for Cancer Genomics (https://www.cbioportal.org) was consulted to look for CDC42 mutations in cancer patients.

### Protein localization and fluorescence microscopy

The localization of GFP-Cdc42p (PC6454) and the GFP-Cdc42p alleles where the expression of the fused protein is driven by the *CDC42* promoter. Wild-type and mutant strains were grown in SD media lacking uracil for plasmid selection for 16h at 30°C, resuspended in fresh media and grown for 4 or 5h to mid-log phase, unless otherwise indicated. To visualize GFP-Cdc42p in the temperature sensitive mutant *cim3-1*, cells were grown for another 2 h at 37°C.

Differential interference contrast (DIC) and fluorescence microscopy using fluorescein isothiocyanate (FITC) and Rhodamine filter sets were performed using an Axioplan 2 fluorescence microscope (Zeiss) with a Paln-Apochromat 100x/1.4 (oil) objective with the Axiocam MRm camera (Zeiss). Images were analyzed using Axiovision 4.4 software (Zeiss). Staining with the lipophilic dye FM4-64 were performed as described in (Amberg et al., 2006). Images were analyzed with ImageJ (https://imagej.nih.gov) and Adobe Photoshop. Fluorescence images are shown in green (GFP) or red (FM4-64) or converted to grayscale and inverted using ImageJ. Confocal microscopy was performed as previously described (Prabhakar et al., 2020).

### Confocal microscopy

Experiments were performed based on (Prabhakar et al., 2020).The *cim3-1*mutant (PC5852) expressing GFP-Cdc42p (PC6454) or GFP-Cdc42p^K16R^ (PC7697) was grown at 30°C for 16 h on SD-URA. 10 μL of cells diluted to 0.02 O.D. were placed under 1% agarose pads using a 12 mm Nunc glass base dish (Cat#150680, Thermo Scientific, Waltham). Cells were grown at 30°C for 3 h prior to imaging. To prevent dehydration, a cotton pad was placed around the agar. Live-cell microscopy was performed with a Zeiss 170 confocal microscope equipped with a Plan-Apochromat 40x/1.4 Oil DIC M27 objective. For the detection of GFP-Cdc42p a 488nm laser (496nm-548nm filter), was used. Images were taken with multiple Z-stacks (8-10) and a distance of 1 μm between each Z-stack. Images were analyzed with ImageJ using the Z-project and template matching plugins.

### Mother Enrichment Program

Mother Enrichment Program cells (Lindstrom and Gottschling, 2009) (PC7713) expressing GFP-Cdc42p (PC6454) were grown 16 h in SD-URA media supplemented with 0.02 mg/l of adenine. 0.005OD (600nm) of cells were resuspended in 10 ml of fresh SD-URA media supplemented with 0.02 mg/l adenine and 1µM of β-estradiol and grown for 48 h at 30°C. Cells were collected using a 0.2 µm pore centrifuge filter and a soft spin (1,000 rpm). Cells were washed twice with phosphate-buffered saline solution (PBS; 137 mM NaCl, 2.7 mM KCl, 10 mM Na_2_HPO_4_, 1.8 mM KH_2_PO_4_) in the column and stained with 20 µg/ml of WGA-conjugates fluorochrome (Wheat Germ Agglutinin, Alexa Fluor 647 Conjugate; Cat# W32466, Thermo Scientific, Waltham) for 30 min on ice. Cells were washed twice with PBS and resuspended into 50 µL of fresh media prior imaging. Cell microscopy was performed with inverted Leica DMI8 THUNDER microscope equipped with HC PL APO 63x/1.20 W CORR CS2 objective. For the detection of GFP-Cdc42p a 490 nm (BP475-BP515 nm), and for WGA-alexa-647 a 660 nm (BP640-BP720 nm) LED filters were used.

### Statistical Analysis

Statistical evaluations were performed with Prism 7 (GraphPad; https://www.graphpad.com/scientific-software/prism/). One-way ANOVA test followed by Tukey’s multiple comparison test were used to compare more than two datasets. Mann-Whitney U test non-parametric test was used to analyzed levels of GFP-Cdc42p at the plasma membrane. Tests were indicated for each experiment in the corresponding figure legend.

## MOVIES

**Movie 1.**
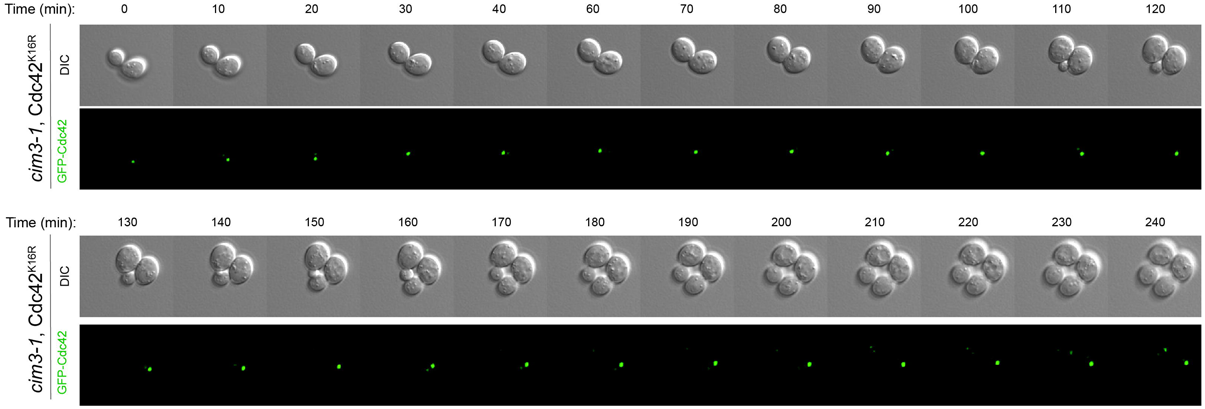
Confocal time-lapse microscopy of the *cim3-1* mutant (PC5852) expressing GFP-Cdc42p (PC6454) on SD-URA media incubated at 37°C. Cells were pre-incubated for 15 min at 37°C before the analysis. Time interval, 10 min.

**Movie 2.**
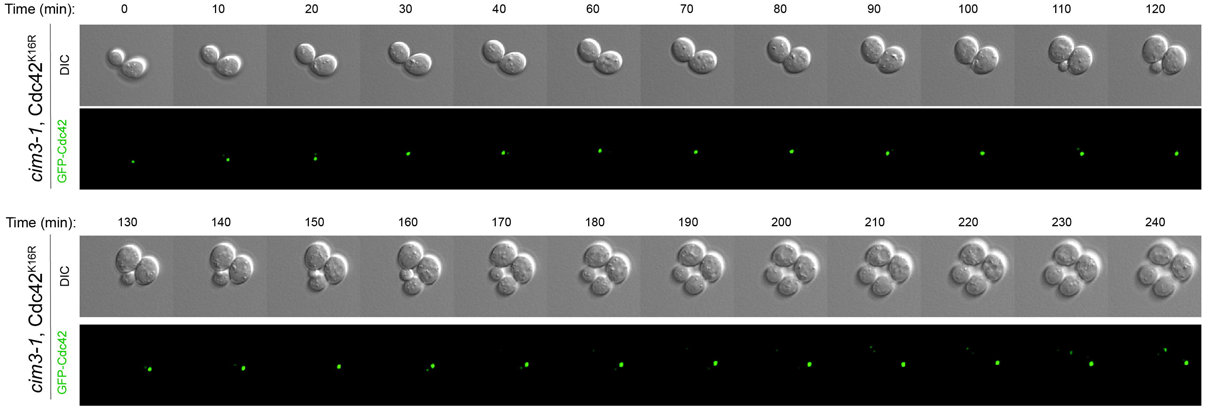
Confocal time-lapse microscopy of the *cim3-1* mutant (PC5852) expressing GFP-Cdc42p^K16R^ (PC7697) on SD-URA media incubated at 37°C. Cells were pre-incubated for 15 min at 37°C before the analysis. Time interval, 10 min.

