## Supplementary figures and images for "Exploring the Regulation of Cdc42 Stability and Turnover in Yeast"

### Figure S1. GFP-Cdc42p accumulated high molecular weight products in a Rsp5-dependent manner

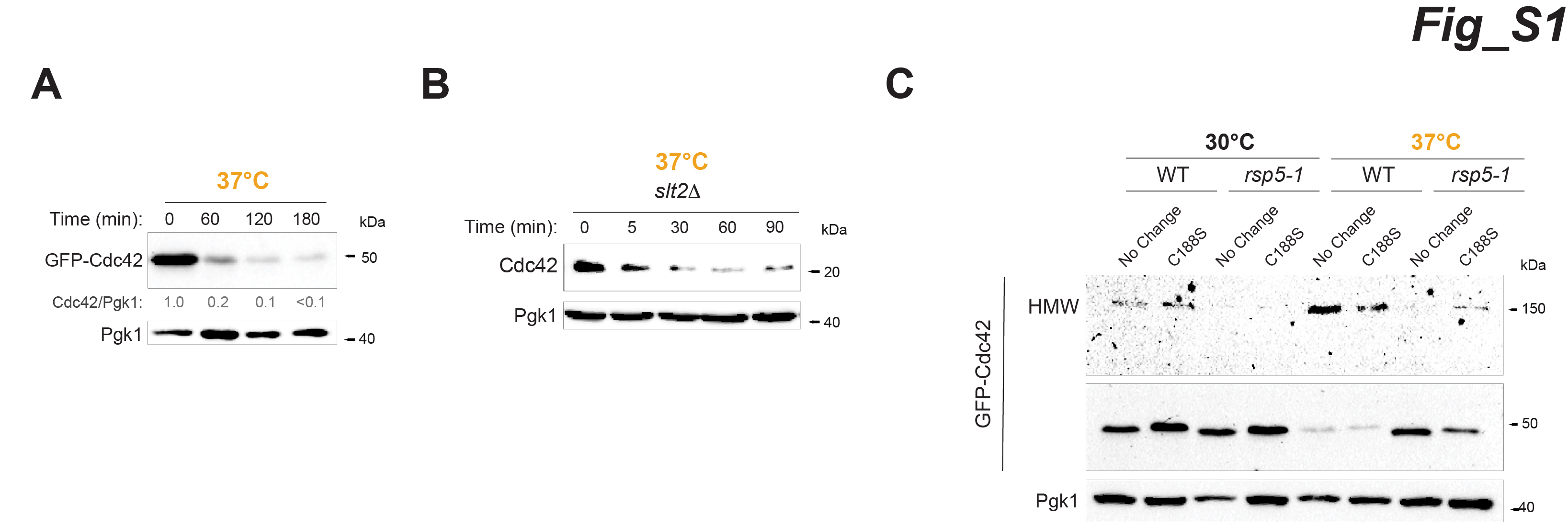

### Figure S3. Role of ESCRT-vacuole pathway in degrading Cdc42p

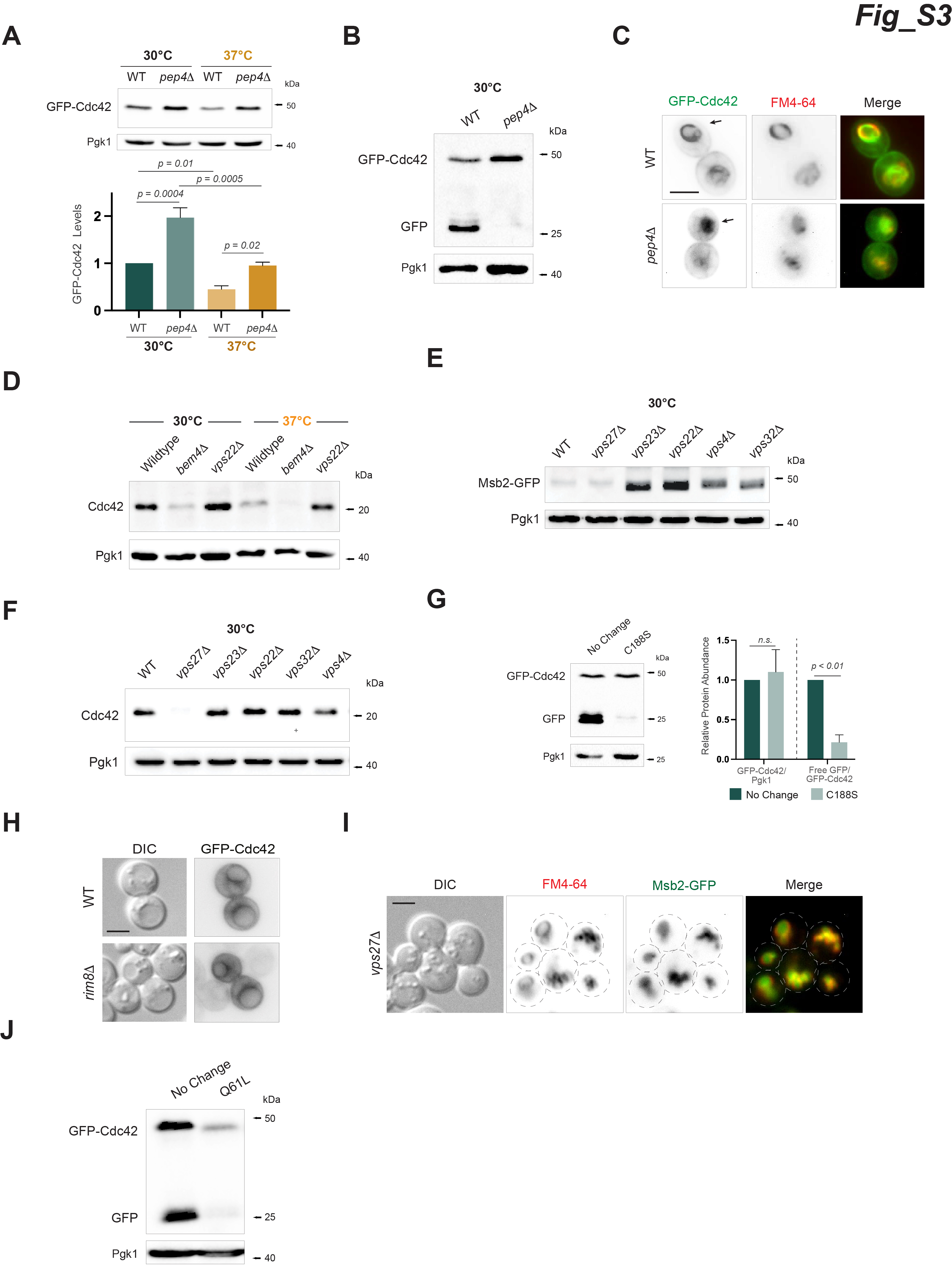

### Figure S5. Cdc42p aggregate formation of lysine mutants.

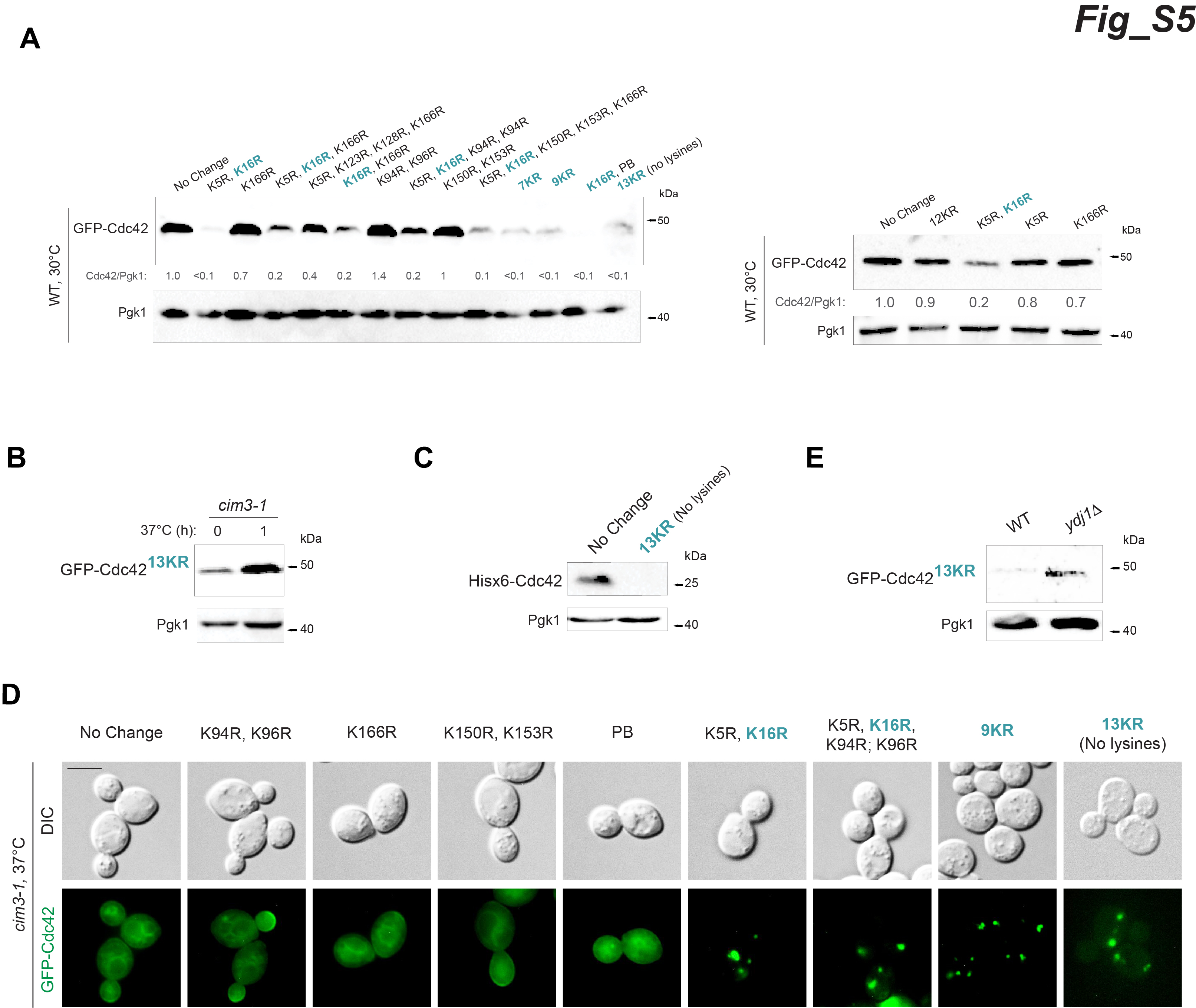

### Figure S6. Overexpression of Cdc42p caused aggregate formation.

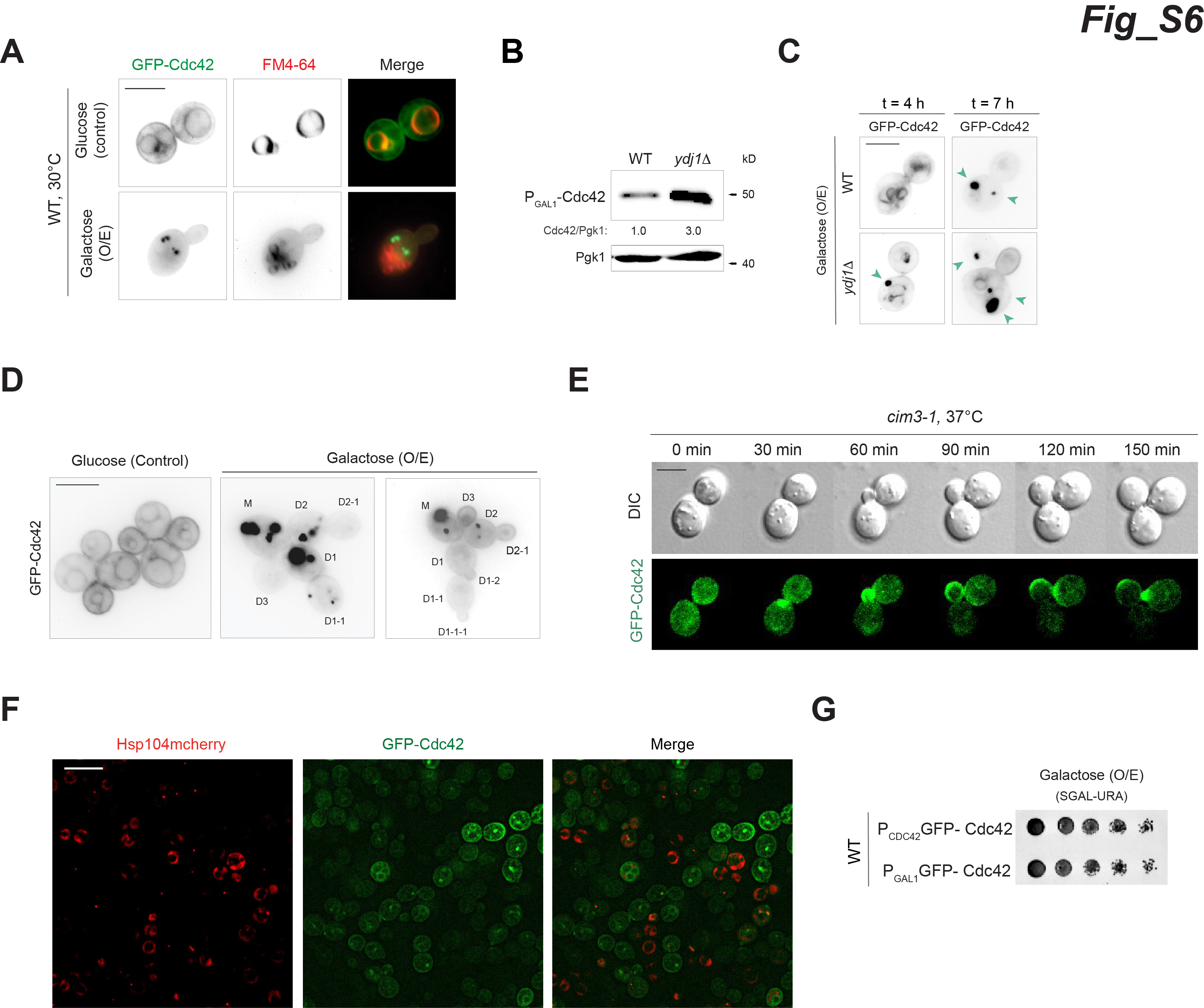

### Supplemental Data 1

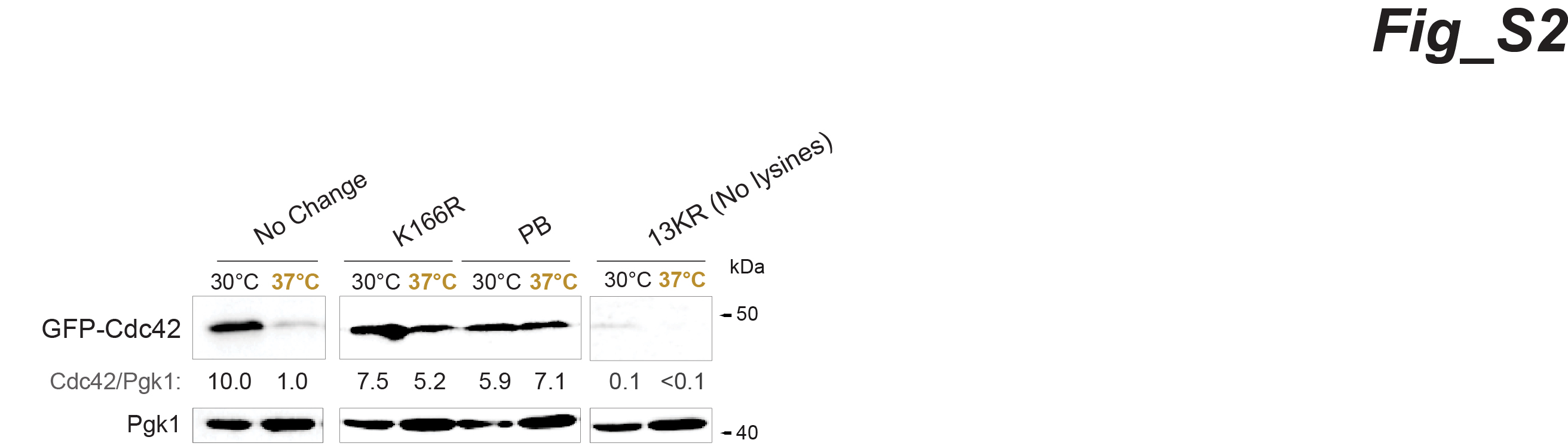

### Supplemental Data 2

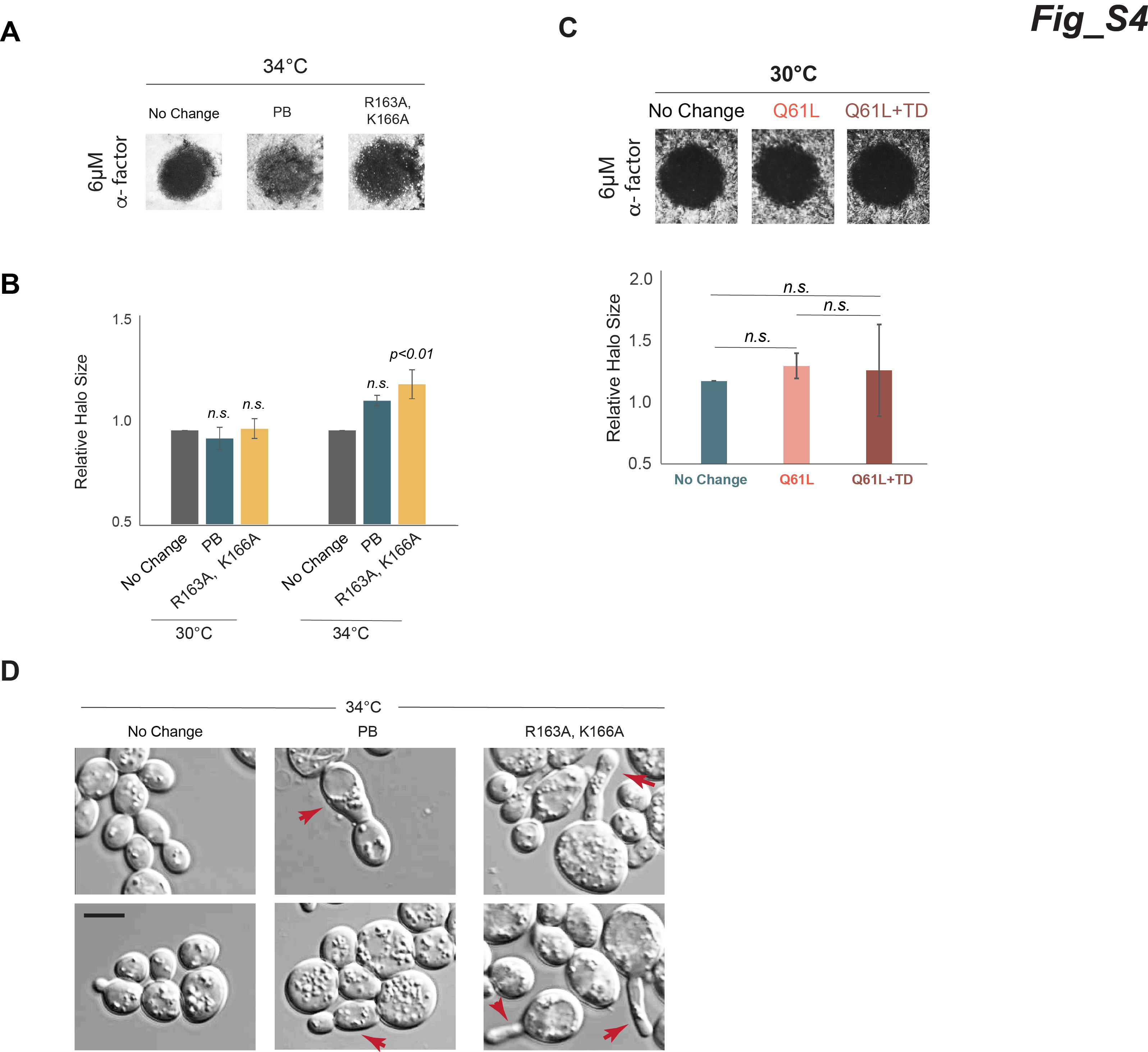
