## Supplemental figure legends for "Exploring the Regulation of Cdc42 Stability and Turnover in Yeast"

Corresponding author: † Paul J. Cullen

532 Cooke Hall

Department of Biological Sciences

State University of New York at Buffalo

Buffalo, NY 14260-1300

### SUPPLEMENTAL FIGURES LEGENDS

**Figure S1. GFP-Cdc42p accumulated high molecular weight products in a Rsp5-dependent manner. A)** Wild-type cells expressing GFP-Cdc42p (PC6454) were grown at 30ºC for 5 h and shifted to 37ºC, samples were analyzed at the indicated time points. Anti-GFP and anti-Pgk1 antibodies were used. **B)** Cells lacking Slt2p were grown at 30ºC for 5 h and shifted to 37ºC for the indicated time points. See *Fig. S1A* for details. **C)** Levels of GFP-Cdc42p (PC6454) or GFP-Cdc42p$^{C188S}$ (PC7350) expressed in wild-type (WT, PC3288) and *rsp5-1* (PC3290) cells incubated 5 h at 30ºC (30ºC) and 2h at 37ºC (37ºC). Top blot corresponds to a longer exposure time than the middle blot. See Fig. S1A for details.

**Figure S2. Lysine residues required for Cdc42p turnover at 37ºC. A)** Wild-type cells (WT; PC538) expressing GFP-Cdc42p (PC6454), GFP-Cdc42p$^{K166R}$ (PC$), GFP-Cdc42p$^{PB}$ (K183R,K184R,K186R,K187R; PC7320), or 13KR (PC7633; GFP-Cdc42$^{K5R, K16R, K94R, K96R, K123R, K128R, K150R, K153R, K166R, K183R, K184R, K186R, K187R}$) were grown at 30ºC for 5 h and shifted to 37ºC for 2 h. See Fig. S1A for details.

**Figure S3. Role of ESCRT-vacuole pathway in degrading Cdc42p. A)** Wild-type cells (PC986, S288C) and the *pep4*Δ (PC3063) mutant expressing GFP-Cdc42p grown at 30ºC for 5 h (30ºC) and shifted to 37ºC for 2 h (37ºC). See Fig. 1D for details. **B)** Wild-type cells (WT, PC986) and the *pep4*Δ (PC3063) mutant expressing GFP-Cdc42p were grown at 30ºC for 5 h. See Fig. S1A for details. **C)** Fluorescence microscopy of wild-type cells (S288C, PC986) and the *pep4*Δ (PC3063) mutant expressing GFP-Cdc42p grown for 5 h at 30ºC and stained with the lipophilic dye FM4-64. Bar 5µm. **D)** Levels of Cdc42p in wild-type cells and *bem4*Δ and *vps22*Δ mutants grown for 5 h at 30ºC (30ºC) and 2 h at 37ºC (37ºC). See Fig. S1A for details. **E)** Levels of Msb2-GFP in wild-type cells and indicated ESCRT mutants grown at 30ºC for 5 h. See Fig. S1A for details. **F)** Levels of GFP-Cdc42p in wild-type cells and indicated ESCRT mutants grown at 30ºC for 5 h. See Fig. S1A for details. **G)** Right, immunoblots analysis of wild-type cells (Σ1278b, PC538) expressing GFP-Cdc42p and GFP-Cdc42p$^{C188S}$. Left, GFP-Cdc42p refers to GFP-Cdc42p levels compared to Pgk1p from two biological replicates. GFP refers to free GFP divided by GFP-Cdc42p intensity from two biological replicates. Error bars represent the standard deviation, and t-test was used to generate p-values. **H)** GFP-Cdc42p localization in wild-type cells (WT) and cells lacking Rim8p (*rim8*Δ). Bar, 5µm. **I)** Fluorescence microscopy of cells lacking Vps27p and expressing Msb2-GFP grown for 5 h at 30ºC and stained with FM4-64 dye. Bar, 5µm. **J)** Wild-type cells (WT, PC538) expressing GFP-Cdc42p or GFP-Cdc42p$^{Q61L}$ were grown at 30ºC for 5 h.

**Figure S4. Morphological defects of turnover-defective versions of Cdc42p at 37ºC. A)** Halo formation in response to α-factor of wild-type (WT, Σ1278b, PC6810) cells and indicated *CDC42* alleles. Cells grown for 16 h were spread on YEPD media where 6 µM α-factor was spotted on the top to study cell cycle arrest, plates were incubated for 48 h at the indicated temperature. Wild-type cells, Cdc42p$^{PB}$, and Cdc42p$^{K163R,K166R}$ alleles were grown on YPD for 48h at 34ºC. **B)** Relative halo size of same cells examined in panel 3C. Data were analyzed by one-way ANOVA followed by a Tukey's multiple comparison test, n= 4, Error bars refer to standard deviation. **C)** Halo formation in WT cells expressing GFP-Cdc42p (Cdc42), GFP-Cdc42p$^{Q61L}$ (Q61L), and GFP-Cdc42p$^{K5R;\ Q61L;\ K94R;\ K96R}$ (Q61L+TD) in response to two concentrations of a-factor, 1.6 µM and 6 µM. Cells were incubated for 48 h at 30ºC. Bottom, data were analyzed by one-way ANOVA followed by a Tukey's multiple comparison test, n= 4, Error bars refer to standard deviation. **D)** Serial dilutions of wild-type cells and *CDC42* alleles grown at 30ºC and 34ºC for 3 days. Bar, 5µm.

**Figure S5. Cdc42p aggregate formation of lysine mutants. A)** Wild-type cells expressing GFP-Cdc42p (No change, PC6454), GFP-Cdc42p$^{K5R,K16R}$ (PC7504), GFP-Cdc42p$^{K166R}$ (PC7518), GFP-Cdc42p$^{K5R,K16R,K166R}$ (PC7506), GFP-Cdc42p$^{K5R,K123R,K128R,K166R}$ (PC7635), GFP-Cdc42p$^{K16R,K166R}$ (PC7506), GFP-Cdc42p$^{K94R,K96R}$ (PC7508), GFP-Cdc42p$^{K5R,K16R,K94R,K96R}$ (PC7507), GFP-Cdc42p$^{K150R,K153R}$ (PC7511), GFP-Cdc42p$^{K5R,K16R,K150R,K153R,K166R}$ (PC7512), GFP-

Cdc42p$^{K5R,K16R,K94R,K96R,K123R,K128R,K166R}$ (7KR; PC7509), and GFP-Cdc42p$^{K5R,K16R,K94R,K96R,K123R,K128R,K150R,K153R,K166R}$ (9KR; PC7513), Cdc42p $^{K16R,K183R,K184R,K186R,K187R}$ (PC7520), and GFP-Cdc42p$^{K5R,K16R,K94R,K96R,K123R,K128R,K150R,K153R,K166R,K183R,K184R,K186R,K187R}$ (13KR; PC7521) grown to mid-log phase at 30ºC. See Fig S1A for details. **B)** *cim3-1* mutant cells expressing Hisx6-Cdc42 (No Change; PC7571) and Hisx6-Cdc42 $^{K5R,K16R,K94R,K96R,K123R,K128R,K150R,K153R,K166R,K183R,K184R,K186R,K187R}$ (13KR; PC7572) grown to mid-log phase at 30ºC. See Fig S1A for details. **C)** Wild-type cells expressing Hisx6-Cdc42 (No change; PC7571) and Hisx6-Cdc42 $^{K5R,K16R,K94R,K96R,K123R,K128R,K150R,K153R,K166R,K183R,K184R,K186R,K187R}$ (13KR; PC7572) grown to mid-log phase at 30ºC. See Fig S1A for details. **D)** Localization of same GFP-Cdc42p alleles indicated in panel 2A expressed in *cim3-1* mutant (PC5852) cells incubated at 37ºC for 2h. Bar, 5µm. **E)** GFP-Cdc42p$^{13KR}$ levels in wild-type cells and the *ydj1*Δ mutant.

**Figure S6. Overexpression of Cdc42p caused aggregate formation. A)** Fluorescence microscopy of Wild-type cells (WT, Σ1278b, PC538) expressing GFP-Cdc42p (control) or pP$_{GAL1}$-GFP-*CDC42* (PC7349) grown for 6 h in YEPD (Glucose) or YEP-GAL (Galactose), respectively and stained with the FM4-64 dye. Bar, 5 µm. **B)** Wild-type (WT) cells and cells lacking Ydj1p expressing pP$_{GAL1}$-GFP-linker-*CDC42P* were grown in YEP-GAL for 3 h and analyzed by fluorescence microscopy. Scale bar, 5 µm. **C)** Same cells as in panel *S6B* were explored by microscopy after 4 h and 7 h grown in YEPGAL media. Bar, 5 µm. **D)** Fluorescence microscopy of wild-type cells expressing pP$_{GAL1}$-GFP-linker-*CDC42P* grown for 8 h in YEP-GAL media. M refers to mother cells, D to daughter and number the order of generation. Bar, 5 µm. **E)** Time-lapse confocal microscopy of the *cim3-1* mutant (PC5852) expressing GFP-Cdc42p (PC6454) grown at 37ºC. Bar, 5µm. **F)** Fluorescence microscopy of Hsp104-mcherry MEP cells expressing GFP-Cdc42p grown for 24h at 30ºC. Bar, 15µm. **G)** Serial dilutions of wild-type cells expressing GFP-Cdc42p ($P_{Cdc42}$GFP-Cdc42) induced by Cdc42p promoter and GAL1-GFP-Cdc42 ($P_{GAL1}$-GFP-Cdc42) induced by the GAL1 promoter on SGAL-URA plates grown for 2 days at 30ºC. E)

### MOVIES

**Movie 1.** Confocal time-lapse microscopy of the *cim3-1* mutant (PC5852) expressing GFP-Cdc42p (PC6454) on SD-URA media incubated at 37ºC. Cells were pre-incubated for 15 min at 37ºC before the analysis. Time interval, 10 min.

**Movie 2.** Confocal time-lapse microscopy of the *cim3-1* mutant (PC5852) expressing GFP-Cdc42p$^{K16R}$ (PC7697) on SD-URA media incubated at 37ºC. Cells were pre-incubated for 15 min at 37ºC before the analysis. Time interval, 10 min.
